# Transcranial ultrasound neuromodulation of a human white matter tract

**DOI:** 10.1101/2025.11.10.687582

**Authors:** Dylan Curtin, Juhi Walinjkar, Elsa F. Fouragnan, Joshua J. Hendrikse, James P. Coxon

**Author notes:** Correspondence and material requests should be addressed to James P. Coxon and Dylan Curtin.

## Abstract

Non-invasive methods to causally modulate human white matter tracts have proved elusive. *In vitro* and *in vivo* studies indicate that myelinated axons are sensitive to mechanical perturbations from ultrasound, providing a mechanistic rationale for targeting white matter tracts with ultrasound neuromodulation. Here, we demonstrate that transcranial ultrasound stimulation (TUS) targeted at the human corticospinal tract induces site-specific changes in motor neurophysiology. In a within-subject study, 15 healthy adults received 5 Hz repetitive TUS targeted at the left corticospinal tract at the level of the superior internal capsule. The homologous tract in the opposite hemisphere and the ipsilateral motor cortex were targeted as comparison conditions to confirm site specificity. Left corticospinal tract TUS increased corticomotor excitability and intracortical facilitation, whereas left motor cortex TUS reduced intracortical inhibition. Supporting analysis using diffusion tractography indicated that the magnitude of these changes depended jointly on ultrasound pressure and white matter microstructure (fractional anisotropy). These findings provide causal evidence that ultrasound can modulate white matter neurophysiology and establish TUS as a tool for non-invasive neuromodulation of white matter tracts in health and disorders of white matter.

**Significance statement:** White matter pathways are central to brain function and disease, yet causal tools to modulate these tracts in humans are lacking. We show that neuronavigated, low-intensity transcranial ultrasound targeted to the corticospinal white matter tract induces site-specific changes in motor neurophysiology. Supporting analyses using diffusion MRI tractography indicated that individual responses depended jointly on ultrasound pressure and local tissue properties, supporting an anatomy-informed, precision approach. These findings establish tract-targeted ultrasound as a non-invasive way to modulate human white matter neurophysiology, opening new opportunities for mechanistic tests of brain circuits and for developing circuit-guided interventions in neurological and psychiatric disorders.

## Introduction

White matter plasticity refers to the experience-dependent changes in myelin and axons that shape neural communication^1–3^. Such changes are critical for sensorimotor, cognitive, and affective functions and support recovery after brain injury^4,5^. Yet, in humans, evidence of white matter plasticity has relied on correlative neuroimaging^1,2^, leaving the causal mechanisms unknown. Brain stimulation offers a path to causal inference^6^, but available techniques are limited. Transcranial magnetic and electrical stimulation lack the depth and spatial precision to selectively engage myelinated tracts, while deep brain stimulation offers precision only through invasive surgery^7^. There is a need for a non-invasive method that can directly and selectively target white matter pathways.

Transcranial ultrasound stimulation (TUS) offers a promising solution^8^. When skull transmission losses are carefully monitored and minimised, ultrasound energy can be non-invasively focused with millimetre precision to modulate both superficial and deep brain structures, a capability not attainable with magnetic or electrical stimulation methods^9^. In humans and non-human primates, repetitive TUS produces network- and regional-level plasticity in cortical and subcortical grey matter^10–19^. For example, an 80-second, 5 Hz TUS protocol^20^ (20 ms ultrasound pulses repeated every 200 ms) targeted at the primary motor cortex and posterior cingulate cortex has been shown to modulate excitatory and inhibitory plasticity for up to 60 minutes after sonication^20–26^. Early clinical work demonstrates the feasibility of targeting deep white matter in patients with treatment-resistant depression^27,28^. What remains untested is whether TUS can selectively modulate a white matter tract in humans, which requires a physiological readout of the sonicated pathway and comparison sites to establish specificity.

Converging experimental^9,29^ and computational^29^ evidence provides a mechanistic rationale for targeting white matter with ultrasound. *In vitro* and *in vivo* models show that low-intensity ultrasound activates mechanosensitive ion channels^30–35^, including Piezo and the two-pore-domain (K2P) potassium channels TREK-1 and TRAAK^36,37^. A notable property of both TREK-1 and TRAAK is that they are highly expressed at nodes of Ranvier^38,39^, where they regulate axon excitability and conduction^36–38,40^, making myelinated axons a plausible substrate for ultrasound neuromodulation. Consistent with this, ultrasound has been shown to evoke action potentials and transiently alter excitability in animal axonal preparations^41,42^, and computational models of peripheral nerve fibres predict that ultrasound can recruit myelinated axons^43^. However, animal nerve preparations differ from human central white matter in geometry, myelination, and ion-channel distribution^44^. Further, in small animals, the compact size of white-matter tracts and their close adjacency to grey matter^45^ make it challenging to disentangle tract-specific effects from collateral activation of grey matter. Whether these findings translate to humans is unclear. Direct evidence is essential to establish whether ultrasound can modulate white matter in humans.

Here, we tested whether TUS targeted at the corticospinal tract (CST) alters motor system neurophysiology. Focusing on the CST allowed us to probe the targeted pathway with transcranial magnetic stimulation (TMS). Individualised tractography quantified ultrasound– tract engagement, and two comparison conditions, the contralateral CST and ipsilateral motor cortex (M1), established specificity. The M1 condition also enabled comparison with prior TUS studies on cortical excitability^20,24,25,46,47^ while introducing refinements in targeting and modelling. We hypothesised that TUS targeted to the left CST would modulate the corticomotor pathway probed by TMS, and that this effect would depend on the site sonicated. Consistent with this, we found that left CST TUS produced a site-specific increase in corticomotor excitability and intracortical facilitation. Further, supporting analysis indicated that the magnitude of these changes depended jointly on ultrasound pressure and white matter microstructure. These results demonstrate site-specific neuromodulation of the human CST, providing causal evidence that ultrasound engages a white matter tract *in vivo*. The findings lay a foundation for mechanistic tests of white matter circuit function and for tract-targeted neuromodulation in neurological and psychiatric disorders.

## Results

Fifteen healthy participants completed a within-subject, crossover design comprising up to four visits (Fig. 1) (see Methods for sample size calculations and sensitivity analysis). Session 1 included MRI for TUS planning and diffusion-weighted imaging. Sessions 2–4 delivered TUS to the left CST, the left M1 hand area, or the contralateral (right) CST. Visits were spaced 1 week apart and scheduled at a consistent time of day. The order of the left CST and left M1 sessions was pre-determined, counterbalanced, and assigned by an independent researcher using a pseudo-randomised schedule. The contralateral right-CST control was scheduled last after completion of the left CST and left M1 sessions.

**Fig. 1.**
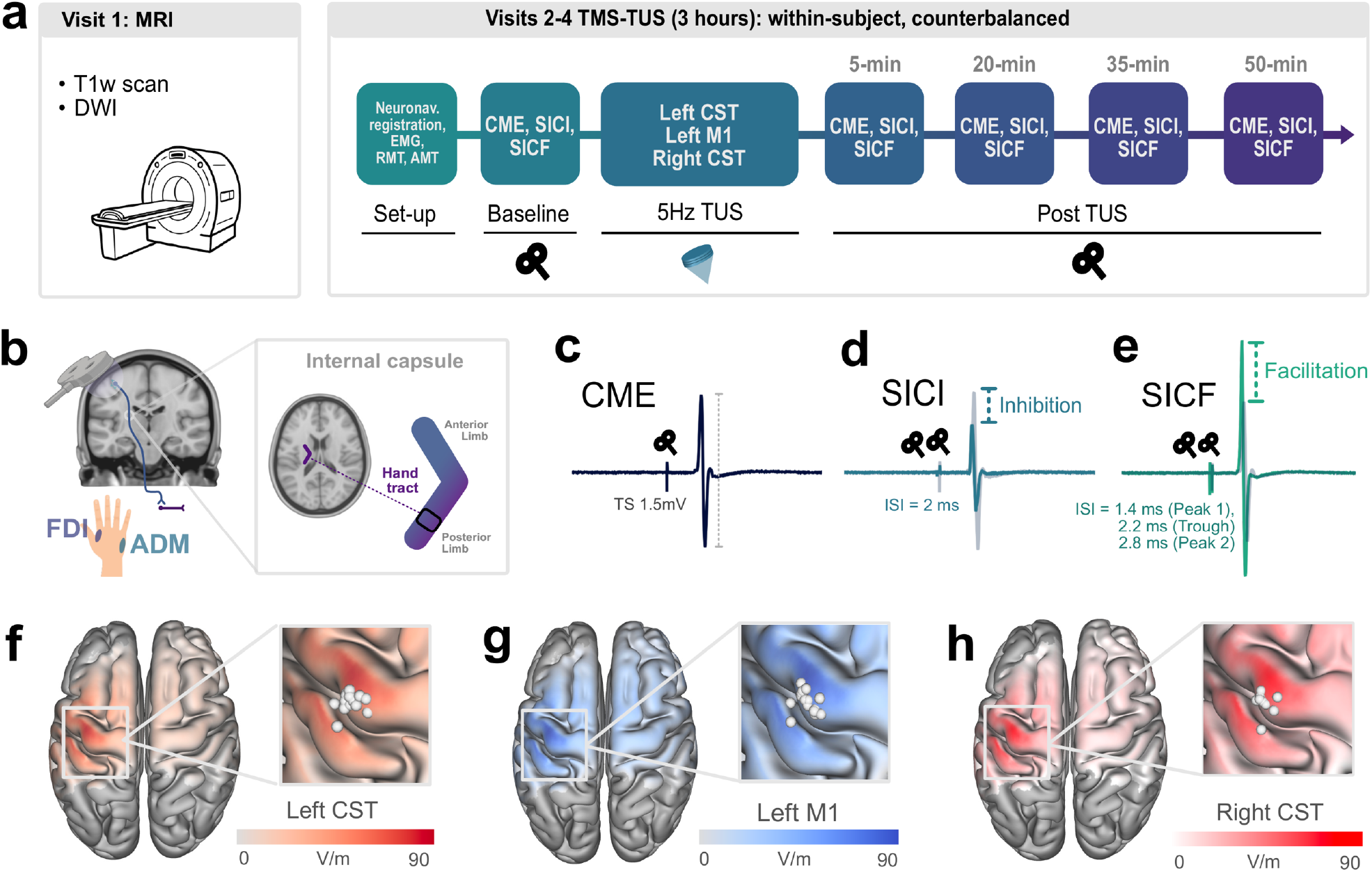
Study design. **a)** Participants completed up to four sessions in a within-subject, crossover design. Session 1 involved anatomical and diffusion-weighted (DWI) MRI scans. In sessions 2–4, baseline measures of excitatory and inhibitory activity were acquired using TMS. Repeat TMS measures were taken 5 minutes after TUS, and repeated at regular intervals. **b)** TMS was optimised to the area of the primary motor cortex corresponding to the first dorsal interosseous (FDI) hand muscle, while also recording EMG from the abductor digiti minimi (ADM). **c**–**e)** Visual depiction of TMS measures. **f**–**h)** Average TMS E-Fields for the left CST, left M1, and right CST sessions in MNI space. White spheres represent the intersection of the TMS vector with the pial surface for each participant. Note: EMG = electromyography, RMT = resting motor threshold, AMT = active motor threshold, CME = corticomotor excitability, SICI = short-interval intracortical inhibition, SICF = short-interval intracortical facilitation.

In each TUS-TMS session, we measured single-pulse corticomotor excitability (CME) and paired-pulse short-interval intracortical inhibition (SICI) and short-interval intracortical facilitation (SICF). For CME, a suprathreshold test stimulus was adjusted to elicit a stable 1.5 mV motor evoked potential (MEP) in the first dorsal interosseous (FDI) at baseline. Recordings from the abductor digiti minimi (ADM) were taken in parallel to assess muscle specificity. For SICI, a subthreshold conditioning pulse preceded the same test pulse by 2 ms. The conditioning intensity was titrated to ∼50% inhibition of the unconditioned MEP. For SICF, the same suprathreshold stimulus was followed by a second stimulus at 100% resting motor threshold (RMT). Inter-stimulus intervals (ISIs) of 1.4, 2.2, and 2.8 ms were included, which correspond to the first peak, trough, and second peak of the SICF time course^24^. All five measures (CME, SICI, SICF_1.4 ms_, SICF_2.2 ms_, SICF_2.8 ms_) were obtained at baseline and repeated at 5, 20, 35, and 50 minutes following sonication.

### Transcranial ultrasound stimulation (TUS)

Fig. 2 summarises the TUS protocol and targeting. Repetitive TUS consisted of a 5 Hz train of pulses (20 ms every 200 ms; 10% duty cycle) at a fundamental frequency of 500 kHz, delivered for 80 s at a spatial-peak pulse-average intensity in water of 20 W/cm² (free field, before skull transmission). We validated the transducer output with in-house hydrophone measurements (Methods). White-noise masking was delivered via bone-conduction headphones in all sessions.

**Fig. 2.**
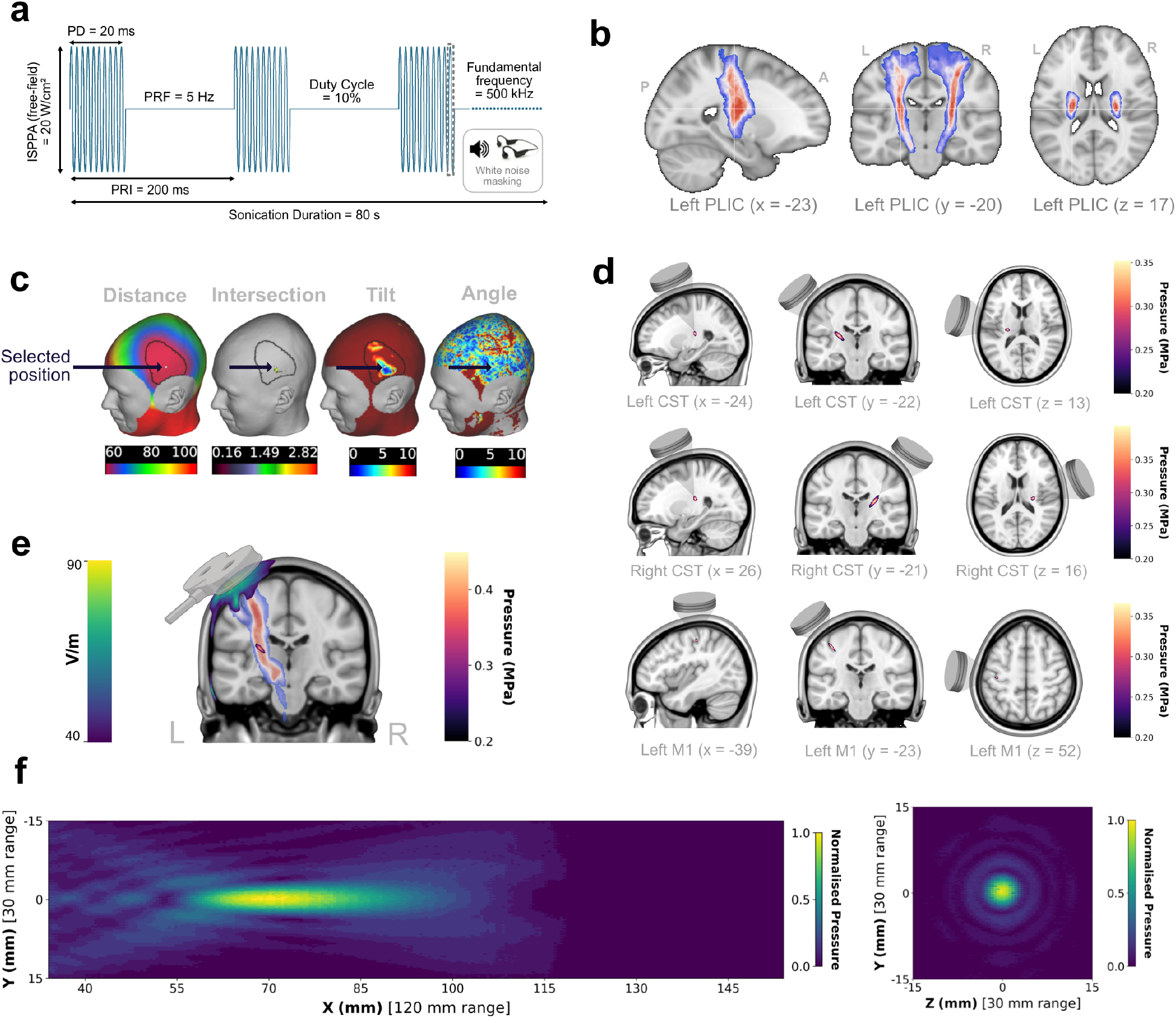
Protocol, targeting, and validation of transcranial ultrasound stimulation (TUS). **a)** 5 Hz repetitive protocol (20 ms pulses every 200 ms; 10% duty cycle) delivered for 80 s with white-noise masking via bone-conduction headphones. **b)** CST targets were defined from a 100% probability CST voxel in the Jülich Histological Atlas, here shown for the left posterior limb of the internal capsule (PLIC). **c)** *PlanTUS* was used to optimise transducer placement for each participant. **d)** Group-averaged simulated transcranial peak pressure maps (MPa) in MNI space for left CST and right CST. Because left M1 targeting was individualised to each participant’s functional anatomy (not an MNI coordinate), we show a representative M1 target. **e)** Individualised probabilistic tractography was used to characterise CST targeting: streamlines are shown seeded from the TMS electric field in cortical grey matter voxels (V/m; left colour bar) with the left CST ultrasound pressure field overlaid (MPa; right colour bar). From the TMS E-field seed region, tracts were retained that passed through a waypoint at the PLIC and terminated in the cerebral peduncle. **f)** Free-field hydrophone measurements (XY and YZ planes) obtained upon completion of data collection validated the focused output of the four-element transducer. *Note*. ISPPA = spatial-peak pulse-average intensity, PD = pulse duration, PRF = pulse repetition frequency, PRI = pulse repetition interval.

Left and right corticospinal tract (CST) targets were defined from 100% probability maps in the Jülich Histological Atlas (posterior limb of the internal capsule, PLIC). Atlas MNI coordinates were nonlinearly inverse warped with FSL’s *FNIRT* to each participant’s T1-weighted image in native space. Transducer placement and beam orientation were optimised with *PlanTUS*^48^ (https://github.com/mlueckel/PlanTUS), and subject-specific acoustic simulations were run to plan the sonication, estimate on-target pressure, and confirm compliance with safety limits^49^. Individualised probabilistic tractography (FSL’s *probtrackx2*) was performed after data collection to quantify target engagement (Fig. 2e). The primary motor cortex (M1) target was localised from hand-knob anatomy and confirmed by TMS ‘hot-spotting’ and motor mapping. Group-averaged ultrasound acoustic pressure maps in MNI space are shown in Fig. 2d, while participant-level maps for each session are provided in Supplementary Figs. S1–S3.

### TMS and TUS characteristics

TMS measures were similar at baseline across sessions (Table 1). For both SICI and SICF, linear mixed effects models confirmed the presence of inhibition and facilitation at baseline, indicated by a main effect of TMS pulse (conditioned, non-conditioned) (SICI: *F*_1,71_ = 35.90, *p* < 0.001; SICF: *F*_1,152_ = 33.24, *p* < 0.001). Both SICF peaks elicited greater facilitation than the SICF trough, β = 1.20, *p* < 0.001 and β = 0.78, *p* < 0.001, respectively. Simulated in-brain ultrasound metrics across the CST and M1 targets are shown in Table 1 (see also Fig. S4).

**Table 1.**
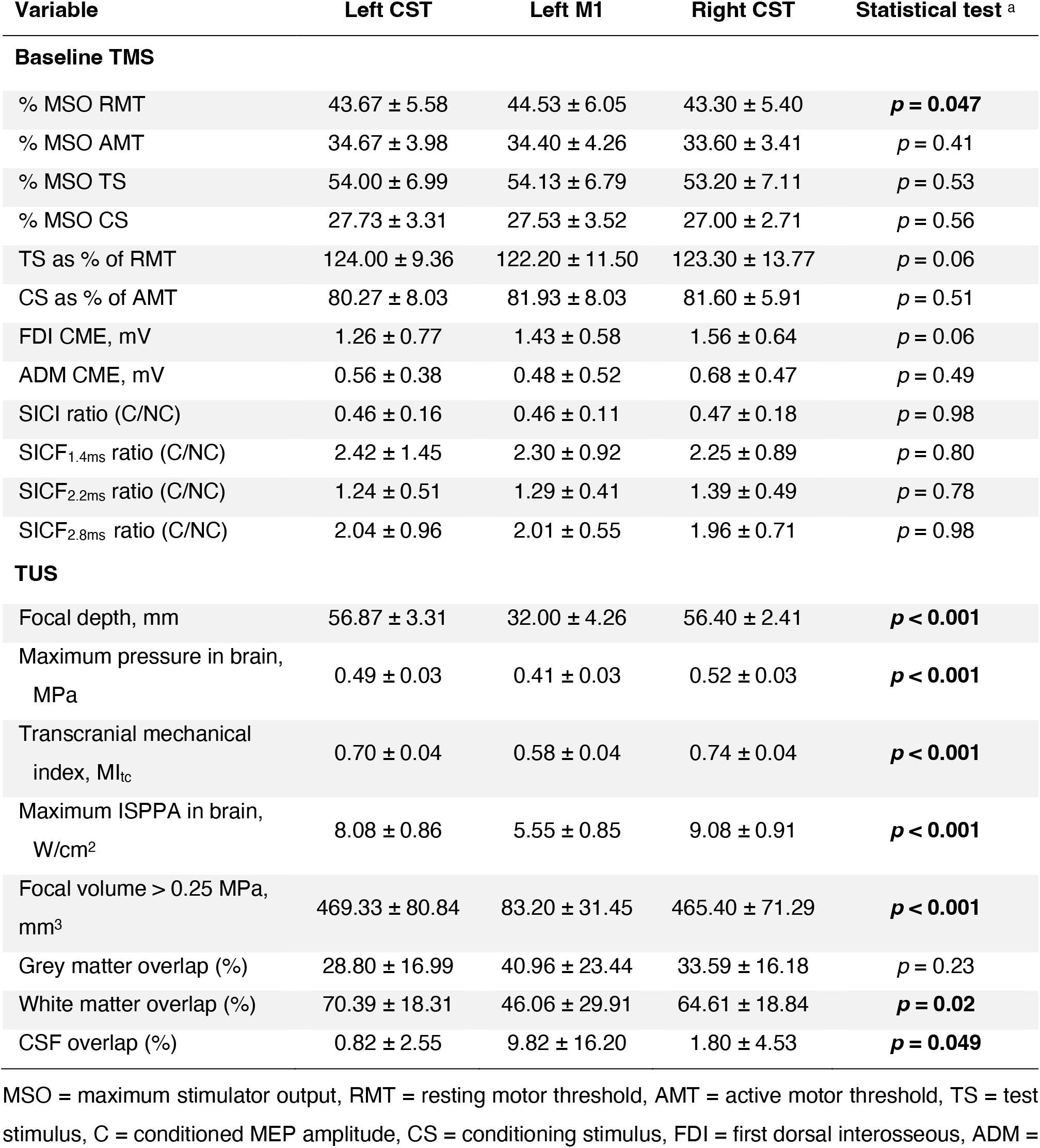
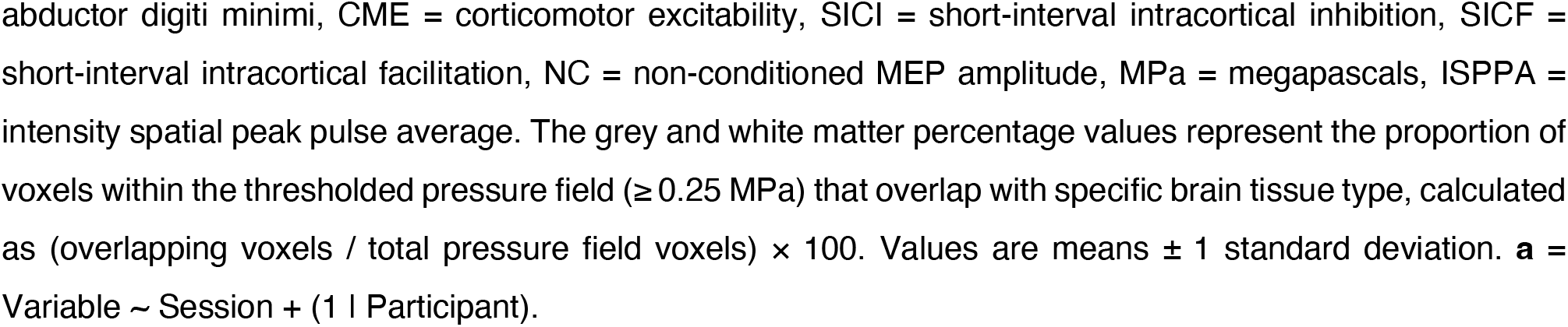
Transcranial magnetic stimulation (TMS) and transcranial ultrasound stimulation (TUS) characteristics by session.

| Variable | Left CST | Left M1 | Right CST | Statistical test <sup>a</sup> |
| --- | --- | --- | --- | --- |
| **Baseline TMS** |  |  |  |  |
| % MSO RMT | 43.67 ± 5.58 | 44.53 ± 6.05 | 43.30 ± 5.40 | **$p = 0.047$** |
| % MSO AMT | 34.67 ± 3.98 | 34.40 ± 4.26 | 33.60 ± 3.41 | $p = 0.41$ |
| % MSO TS | 54.00 ± 6.99 | 54.13 ± 6.79 | 53.20 ± 7.11 | $p = 0.53$ |
| % MSO CS | 27.73 ± 3.31 | 27.53 ± 3.52 | 27.00 ± 2.71 | $p = 0.56$ |
| TS as % of RMT | 124.00 ± 9.36 | 122.20 ± 11.50 | 123.30 ± 13.77 | $p = 0.06$ |
| CS as % of AMT | 80.27 ± 8.03 | 81.93 ± 8.03 | 81.60 ± 5.91 | $p = 0.51$ |
| FDI CME, mV | 1.26 ± 0.77 | 1.43 ± 0.58 | 1.56 ± 0.64 | $p = 0.06$ |
| ADM CME, mV | 0.56 ± 0.38 | 0.48 ± 0.52 | 0.68 ± 0.47 | $p = 0.49$ |
| SICI ratio (C/NC) | 0.46 ± 0.16 | 0.46 ± 0.11 | 0.47 ± 0.18 | $p = 0.98$ |
| SICF<sub>1.4ms</sub> ratio (C/NC) | 2.42 ± 1.45 | 2.30 ± 0.92 | 2.25 ± 0.89 | $p = 0.80$ |
| SICF<sub>2.2ms</sub> ratio (C/NC) | 1.24 ± 0.51 | 1.29 ± 0.41 | 1.39 ± 0.49 | $p = 0.78$ |
| SICF<sub>2.8ms</sub> ratio (C/NC) | 2.04 ± 0.96 | 2.01 ± 0.55 | 1.96 ± 0.71 | $p = 0.98$ |
| **TUS** |  |  |  |  |
| Focal depth, mm | 56.87 ± 3.31 | 32.00 ± 4.26 | 56.40 ± 2.41 | **$p < 0.001$** |
| Maximum pressure in brain, MPa | 0.49 ± 0.03 | 0.41 ± 0.03 | 0.52 ± 0.03 | **$p < 0.001$** |
| Transcranial mechanical index, MI<sub>tc</sub> | 0.70 ± 0.04 | 0.58 ± 0.04 | 0.74 ± 0.04 | **$p < 0.001$** |
| Maximum ISPPA in brain, W/cm<sup>2</sup> | 8.08 ± 0.86 | 5.55 ± 0.85 | 9.08 ± 0.91 | **$p < 0.001$** |
| Focal volume > 0.25 MPa, mm<sup>3</sup> | 469.33 ± 80.84 | 83.20 ± 31.45 | 465.40 ± 71.29 | **$p < 0.001$** |
| Grey matter overlap (%) | 28.80 ± 16.99 | 40.96 ± 23.44 | 33.59 ± 16.18 | $p = 0.23$ |
| White matter overlap (%) | 70.39 ± 18.31 | 46.06 ± 29.91 | 64.61 ± 18.84 | **$p = 0.02$** |
| CSF overlap (%) | 0.82 ± 2.55 | 9.82 ± 16.20 | 1.80 ± 4.53 | **$p = 0.049$** |

MSO = maximum stimulator output, RMT = resting motor threshold, AMT = active motor threshold, TS = test stimulus, C = conditioned MEP amplitude, CS = conditioning stimulus, FDI = first dorsal interosseous, ADM =
abductor digiti minimi, CME = corticomotor excitability, SICI = short-interval intracortical inhibition, SICF = short-interval intracortical facilitation, NC = non-conditioned MEP amplitude, MPa = megapascals, ISPPA = intensity spatial peak pulse average. The grey and white matter percentage values represent the proportion of voxels within the thresholded pressure field ( $\geq 0.25$ MPa) that overlap with specific brain tissue type, calculated as (overlapping voxels / total pressure field voxels) $\times 100$ . Values are means $\pm 1$ standard deviation. **a** = Variable ∼ Session + (1 | Participant).

### Corticomotor excitability (CME; Fig. 3a–d)

TUS altered corticomotor excitability (CME) in a site-dependent manner, measured from the FDI representation targeted by TMS (Session × Time interaction *p* = 0.048; Table 2). Linear trend analysis showed that CME increased more over time after TUS to the left CST compared to right CST (β = 3.34, *p* < 0.001, *d* = 0.84) and more than after left M1 (β = 1.69, *p* = 0.045, *d* = 0.53). A time point contrast demonstrated higher CME for left CST than for right CST (*t*₁₆₆ = 3.71, *p* < 0.001, *d* = 0.62) at the 50-minute time point. As a comparison to past work^20,24,25^, we also compared the effects of left M1 with right CST TUS. Although the overall linear trend contrast was not significant (β = 1.65, *p* = 0.08, *d* = 0.54), the 50-minute time point showed higher CME for left M1 than for right CST (*t*₁₆₆ = 2.00, *p* = 0.047, *d* = 0.40). No other FDI time point contrasts reached significance.

**Fig. 3.**
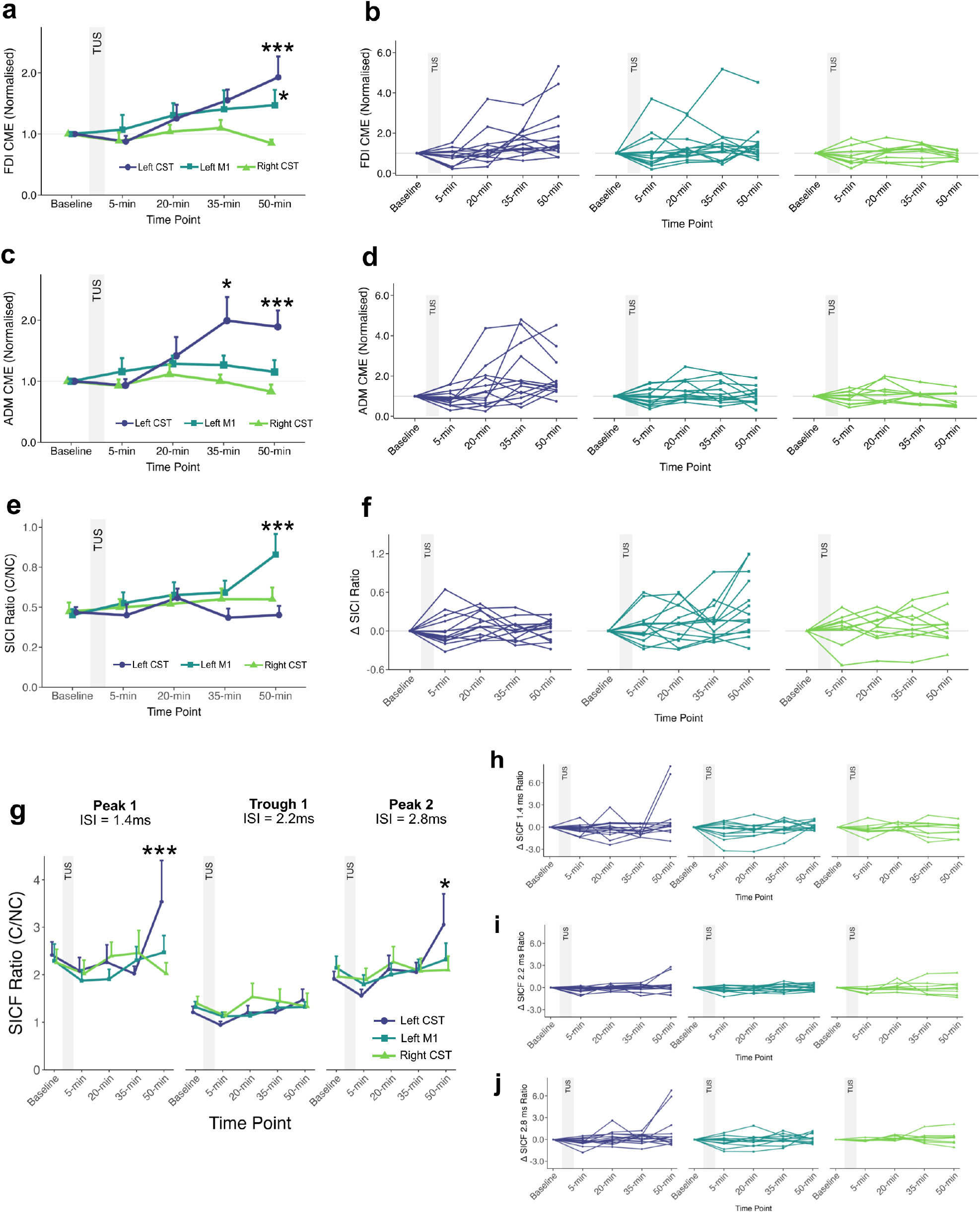
Transcranial ultrasound stimulation modulates motor system excitability and intracortical circuits. Time course of TMS measures across the three TUS conditions: left corticospinal tract (Left CST; *n* = 15), left primary motor cortex (Left M1; *n* = 15), right corticospinal tract (Right CST; *n* = 10). Group-level means (+ 1 s.e.m.) are shown in **a, c, e, and g**. Individual-level data are shown in **b, d, f, h, i, j**. First dorsal interosseous (FDI) and abductor digiti minimi (ADM) motor evoked potential amplitudes (a–d) were normalised to baseline for visualisation (all statistics were performed on non-normalised data). SICI and SICF ratios (e–j) are expressed as a ratio of the conditioned / non-conditioned amplitude. Asterisks indicate significant pairwise differences between TUS sessions at the corresponding time point, \*\*\**p* < 0.001, \*\**p* < 0.01, \**p* < 0.05.

**Table 2.** Transcranial magnetic stimulation (TMS) results.

| Measure | Session | Time | Session × Time |
| --- | --- | --- | --- |
| CME, FDI <sup>a</sup> | $F_{2, 167.70} = 0.93$, $p = 0.40$ | $F_{4, 164.70} = 5.58$, $p < 0.001$ | $F_{8, 164.70} = 2.01$,<br>$p = 0.048$ |
| CME, ADM <sup>b</sup> | $F_{2, 170.39} = 3.72$, $p = 0.026$ | $F_{4, 169.10} = 1.63$, $p = 0.17$ | $F_{8, 169.10} = 2.91$,<br>$p = 0.005$ |
| SICI (C/NC) <sup>c</sup> | $F_{2, 169.54} = 7.17$, $p = 0.001$ | $F_{4, 164.79} = 2.60$, $p = 0.038$ | $F_{8, 164.79} = 2.09$,<br>$p = 0.039$ |
| SICF (C/NC) <sup>d</sup> | $F_{2, 518.02} = 0.17$, $p = 0.84$ | $F_{4, 513.42} = 5.88$, $p < 0.001$ | $F_{8, 513.43} = 3.00$,<br>$p = 0.003$ |

Bolded values denote statistically significant results. CME = corticomotor excitability, FDI = first dorsal interosseous, ADM = abductor digiti minimi, SICI = short-interval intracortical inhibition, SICF = short-interval intracortical facilitation, C = conditioned MEP amplitude, NC = non-conditioned MEP amplitude. Separate linear mixed effects models were constructed for each outcome measure, as follows: **a** = FDI CME ∼ Session × Time + (1 | Participant), **b** = ADM CME ∼ Session × Time + (1 | Participant), **c** = SICI ∼ Session × Time + (1 | Participant), **d** = SICF ∼ Session × Time × Inter Stimulus Interval + (1 | Participant).

Consistent with the FDI results, left CST stimulation had a selective effect on CME recorded from ADM, indicated by a Session × Time interaction (*p* = 0.005; Table 2). The linear increase over time was greater for left CST than for right CST (β = 1.61, *p* = 0.001, *d* = 1.20) and greater than for left M1 (β = 1.31, *p* = 0.002, *d* = 0.90). There was no difference between left M1 and right CST (β = 0.30, *p* = 0.50, *d* = 0.30). At both 35 and 50 minutes, CME was higher for left CST than for right CST (*t*₁₇₀ = 2.58, *p* = 0.011, *d* = 0.52 and *t*₁₇₀ = 3.60, *p* < 0.001, *d* = 0.75, respectively) and higher for left CST than for left M1 (*t*₁₆₉ = 2.52, *p* = 0.013, *d* = 0.46 and *t*₁₆₉ = 3.85, *p* < 0.001, *d* = 0.68, respectively). No other time point comparisons were significant.

### Short-interval intracortical inhibition (SICI; Fig. 3e & f)

In contrast to CME, changes in inhibition were specific to the M1 session (Session × Time interaction; *p* = 0.039; Table 2). The SICI ratio increased after left M1 TUS (i.e. the conditioned response showed reduced suppression relative to the non-conditioned response), indicating a reduction in inhibition, and this differed from left CST (β = 0.87, *p* = 0.002, *d* = 0.66) but not right CST (β = 0.61, *p* = 0.06, *d* = 0.46). The two CST sessions did not differ (β = 0.26, *p* = 0.353, *d* = 0.31). At 50 minutes, there was a greater decrease in inhibition in the left M1 condition compared to the left CST (*t*₁₆₆ = 4.84, *p* < 0.001, *d* = 0.75) and right CST (*t*₁₆₆ = 3.39, *p* = 0.001, *d* = 0.52) conditions. No other time point comparisons were significant.

### Short-interval intracortical facilitation (SICF; Fig. 3g–j)

Consistent with the CME results, SICF varied over time across sessions (Session × Time, *p* = 0.003, Table 2). This interaction is our key SICF result. To characterise this interaction further, we ran an exploratory analysis comparing the left and right CST sessions at each inter-stimulus interval, with contrasts Holm-corrected across the three intervals. At 50 minutes, facilitation after left CST TUS was increased at the 1.4 ms I-wave peak (β = 1.40, *t*₅₁₅ = 3.88, *p* < 0.001, *d* = 0.37) with a smaller increase at the 2.8 ms peak (β = 0.85, *t*₅₁₅ = 2.27, *p* = 0.047, *d* = 0.35), and no change at the 2.2 ms trough (β = 0.02, *t*₅₁₅ = 0.07, *p* = 0.95, *d* = 0.16). This effect was also present in the counterbalanced comparison, with greater facilitation after left CST than left M1 TUS at 50 minutes (β = 0.96, *t*₅₁₄ = 2.91, *p* = 0.004, *d* = 0.31). Linear trend contrasts comparing the left and right CST across the full time course did not survive correction across the three intervals (1.4 ms *p* = 0.13, 2.8 ms *p* = 0.13).

### Probabilistic tractography findings

The physiological effects of left CST-TUS were consistent with engagement of the targeted white matter pathway. To examine this account further, we first tested whether ultrasound coupling was selective to the tract. This analysis indicated that ultrasound pressure was spatially concentrated within the targeted tract. We then conducted two exploratory analyses, asking whether greater tract engagement was associated with larger physiological change, and whether tract microstructure moderated this relationship. These analyses were consistent in direction with the interpretation that ultrasound acted through the intended pathway.

#### Selectivity of tract engagement (Fig. 4a)

In every participant, the peak in-brain pressure fell within the left CST mask. To further quantify spatial specificity, we compared the 99^th^ percentile pressure within three masks: (i) left CST, (ii) the M1 TMS E-field mask (> 40 V/m), and (iii) whole-brain grey matter (Fig. 4a). Paired *t*-tests showed higher pressure in CST than in M1 (*t*₁₃ = 13.54, *p* < 0.001) and adjacent grey matter (*t*₁₃ = 14.84, *p* < 0.001). Simulated pressure deposition for the left CST-TUS session was therefore concentrated within the targeted tract and was greater than at the cortical TMS stimulation site.

**Fig. 4.**
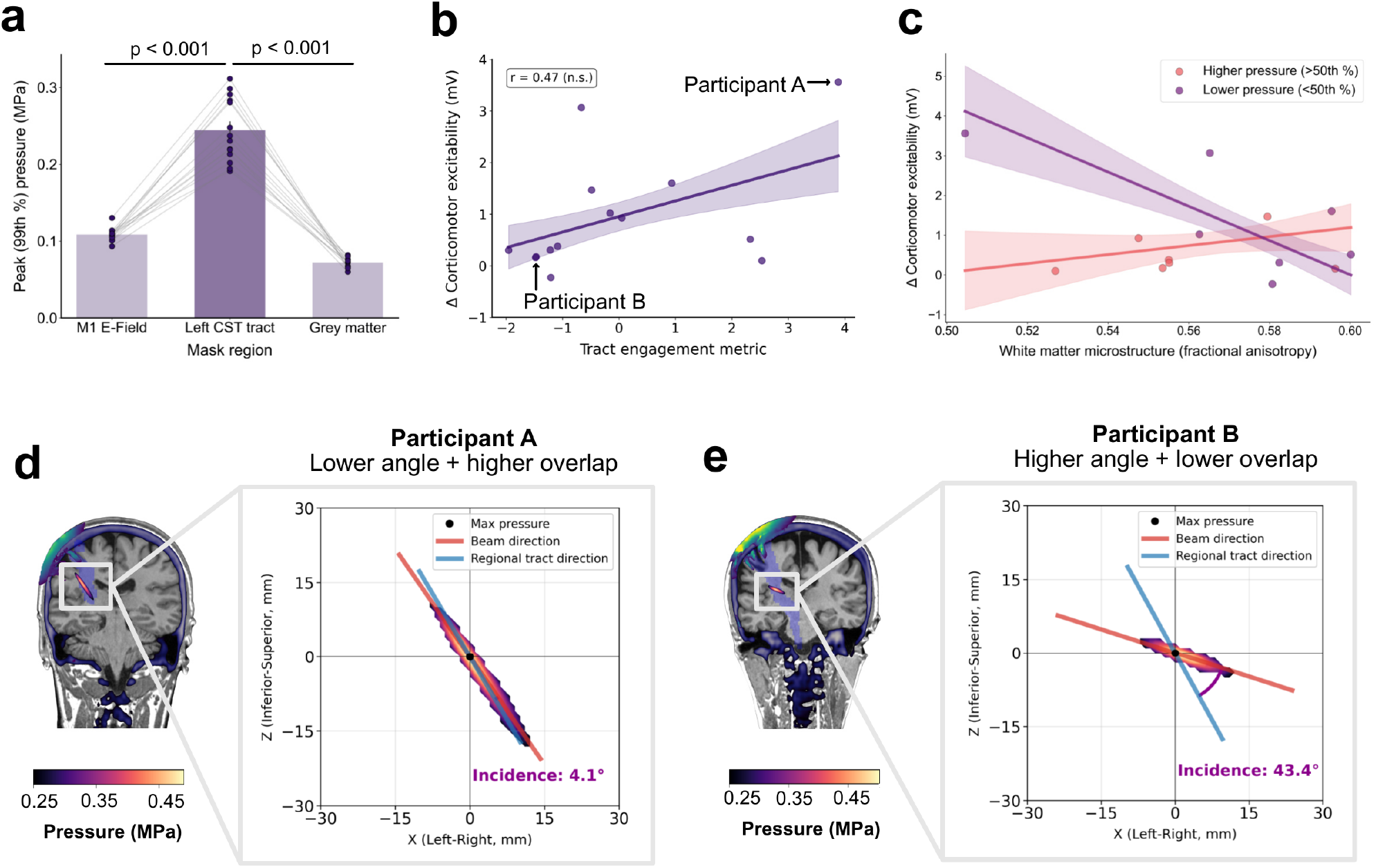
Tractography analyses. **a)** Peak (99^th^ percentile) pressures for the left CST-TUS session were significantly higher in the left CST mask than in the M1 E-field region or adjacent grey matter (both *p* < 0.001, paired *t*-tests). Bars show mean ± s.e.m.; circles indicate individual participants. **b)** Left CST-TUS tract engagement, defined as a composite of CST focal volume overlap and beam–tract alignment, showed a moderate, non-significant positive association with the increase in corticomotor excitability (Pearson’s *r* = 0.47, 95% CI [−0.08, 0.80], *p* = 0.088, two-tailed; Spearman’s ρ = 0.51, *p* = 0.067). Regression line ± s.e.m. shaded. **c)** Fractional anisotropy (FA) moderates the pressure–response relationship (FA × peak simulated in-brain pressure [MPa] interaction: standardised β = 1.36, 95% CI [0.03, 2.70]; *R*² = 0.45; *t*₁₀ = 2.27; p = 0.046). For visualisation, participants are median split by pressure (> 50^th^ percentile = pink; < 50^th^ percentile = purple); lines show fitted regressions with s.e.m. bands. **d,e)** Example participants illustrating lower-incidence angle/higher-overlap targeting and higher-incidence angle/lower-overlap targeting. Vectors show the ultrasound beam (red), the tract direction (blue), and the location of the maximal pressure (black dot). The TMS E-field (V/m) is shown in the *viridis* colour map.

#### Tract engagement index (Fig. 4b, d, e)

We next developed a composite engagement metric incorporating (a) spatial overlap between the tract and the ultrasound focal volume and (b) directional alignment between tract orientation and beam trajectory (where the incidence angle was recoded so that higher values reflected better alignment). Tract engagement showed a moderate, non-significant positive association with the TUS-induced increase in corticomotor excitability (Pearson’s *r* = 0.47, 95% CI [−0.08, 0.80], *p* = 0.088, two-tailed; Spearman’s ρ = 0.51, *p* = 0.067). The direction and magnitude of this association are consistent with an independent report in which target engagement, quantified as overlap between the simulated pressure field and a functionally defined target, predicted the strength of online TUS effects on visual evoked potentials^50^.

#### Individual variability in tract microstructure (Fig. 4c)

Finally, we examined whether local microstructure moderated the pressure–response relation. Mean FA was extracted from a window ± 5 mm along the superior–inferior (Z) axis centred on each participant’s peak-pressure voxel within the left CST. A least-squares model revealed an FA × pressure interaction (standardised β = 1.36, 95% CI [0.03, 2.70], *t*₁₀ = 2.27, *p* = 0.046). The pressure– response slope was more positive in participants with higher FA, indicating that tract organisation moderated the relationship between ultrasound pressure and corticomotor excitability, consistent with more efficient transmission in well-organised tracts. Given the sample size and because simulated pressures carry uncertainty (Methods), we interpret this interaction as supportive evidence.

### Control analyses

We conducted a series of control analyses to confirm the precision and validity of our stimulation procedures. First, we verified the consistency of TMS coil (Fig. 5a) and TUS transducer (Fig. 5b) positioning across sessions. Positioning accuracy was high (mean target error 0.55 ± 0.17 mm for TMS and 0.50 ± 0.26 mm for TUS), with no systematic differences across sessions for either modality (all *p* ≥ 0.07). Second, motor mapping confirmed that the TMS coil location on the scalp (the ‘hotspot’) was optimised for the FDI muscle (Fig. 5c). Finally, we quantified the distance between the TMS coil centre and the TUS transducer centre on the scalp. Prior studies^20,24^ have positioned TUS by placing the transducer directly over the TMS hotspot. As expected, when we quantified distance at the scalp the positions of the TMS coil and TUS transducer showed substantial variability across participants (Fig. 5d), consistent with prior observations that the TMS hotspot is an unreliable proxy for optimal TUS targeting. By contrast, our approach prioritised alignment in the motor cortex: we positioned TUS to overlap the M1 hand region identified in each individual’s anatomy and E-field model (Fig. 1g). For full details of these analyses, see Supplementary Tables S1 and S2, and Supplementary Figs. S1–S3. Verification of EMG signal quality (Fig. S6), control tests to rule out potential order effects, and ultrasound intensity differences are provided in the Supplementary Materials.

**Fig. 5.**
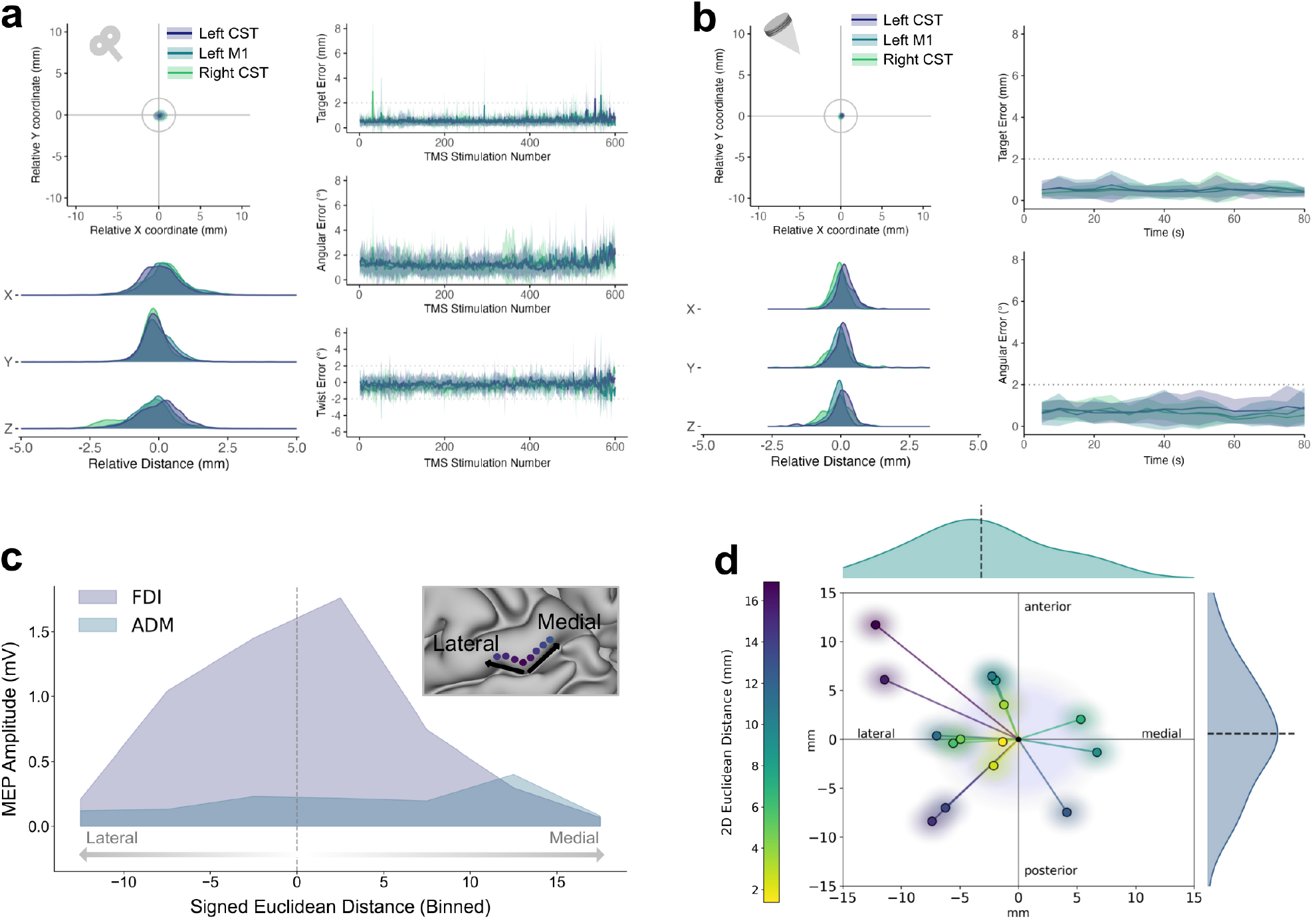
Control analyses. **a)** TMS coil and **b)** TUS transducer vectors were recorded in the neuronavigation software and analysed offline to confirm the accuracy and consistency of TMS and TUS delivery. **c)** A motor mapping procedure verified that the TMS ‘hotspot’ was optimised to the FDI position, whereas the ADM position was located 12.5 mm more medial. **d)** Position of the TUS transducer relative to the TMS coil. The gradient circles are a visual reminder that TMS and TUS stimulate/sonicate a region, not a single point.

## Discussion

Drawing on evidence that ultrasound interacts with myelinated axons, we tested whether transcranial ultrasound stimulation of a white matter tract alters motor system neurophysiology in humans. We demonstrated that TUS of the corticospinal tract (CST) produced a site-specific increase in corticomotor excitability and intracortical facilitation. Acoustic simulations indicated that peak pressures were concentrated within the CST rather than surrounding grey matter or cortical regions, supporting the spatial selectivity of tract engagement. Further, supporting analysis indicated that the magnitude of these effects depended jointly on ultrasound pressure and tract microstructure (fractional anisotropy). Overall, these findings provide causal evidence that ultrasound can exert a neuromodulatory effect on a white matter tract and establish a foundation for precision interventions targeting white matter pathways.

### TUS-induced neuromodulation of a white matter tract

There is accumulating evidence that the neuromodulatory effects of TUS are driven by the activation of mechanosensitive ion channels^30,36,37^, such as TRAAK and TREK-1, which are highly expressed at nodes of Ranvier^38,39^. Ultrasound can also engage glia^51^. In mice, low-intensity ultrasound activates astrocytic TRPA1, driving Ca²⁺ influx and glutamate release that engages NMDA receptors and modulates network excitability^52^. Glia-to-neuron signalling is another plausible route through which sonication could influence axonal conduction and synaptic drive in connected circuits. These findings raise the possibility that myelinated tracts may be sensitive to ultrasound modulation, yet this has not been demonstrated in humans. Our results provide converging evidence that TUS delivered to a white matter tract produces lasting physiological change. First, TMS revealed that left CST-TUS increased corticomotor excitability and intracortical facilitation. Crucially, this effect was site-specific: it was greatest when TUS targeted the same tract probed by TMS, exceeding both the ipsilateral motor cortex and the homologous tract in the opposite hemisphere. Second, in an exploratory analysis, the pressure–response relationship was steeper in participants with higher tract microstructure (fractional anisotropy), indicating that the effect was constrained by the biological and biophysical properties of the targeted tract. Third, acoustic modelling indicated that peak left CST-TUS ultrasound pressures were concentrated in the CST rather than in adjacent grey matter or at the cortical TMS site, consistent with preferential exposure of the targeted tract. We note, however, that approximately 30% of the thresholded field fell within grey matter (Table 1). This is expected given the geometry of the region: the focal volume exceeded the cross-sectional dimensions of the posterior limb of the internal capsule, so the exposed volume necessarily extended into adjacent structures. The tract was therefore the principal but not the only tissue exposed, and a contribution from adjacent grey matter cannot be excluded. Together, these findings provide causal evidence that ultrasound can modulate white matter neurophysiology.

Whether these effects reflect plasticity within the tract itself is an open question. Our TMS measures index the neurophysiological state of the corticomotor pathway as a whole, and cannot localise change to the sonicated white matter. Two accounts remain compatible with our results (Fig. 6). TUS may have produced lasting changes within the CST itself, altering axonal excitability or conduction so that the tract transmits differently^53^. Alternatively, TUS may have transiently altered the efficacy of signal transduction in the CST, with the resulting change driving lasting plasticity at cortical synapses through retrograde (antidromic), or thalamocortical routes, requiring no lasting change in the tract. The first account is consistent with recent evidence that deep brain stimulation of subcallosal cingulate white matter increases fractional anisotropy and myelination in the macaque cingulum bundle^54^. That work differs from ours in stimulation modality and in timescale, since remodelling followed weeks of repeated stimulation rather than a single session. Nonetheless, it supports white matter as a viable substrate for stimulation-induced change. *De novo* myelin synthesis is too slow to contribute to our results^55^, but is not the only form of white matter plasticity, and structural and glial changes have been detected on a much shorter timescale of minutes-to-hours^56–58^. Distinguishing these accounts in humans will require additional measures such as pre-and post-sonication neurite orientation dispersion and density imaging^59^, or diffusion-weighted magnetic resonance spectroscopy^60^.

**Fig. 6.**
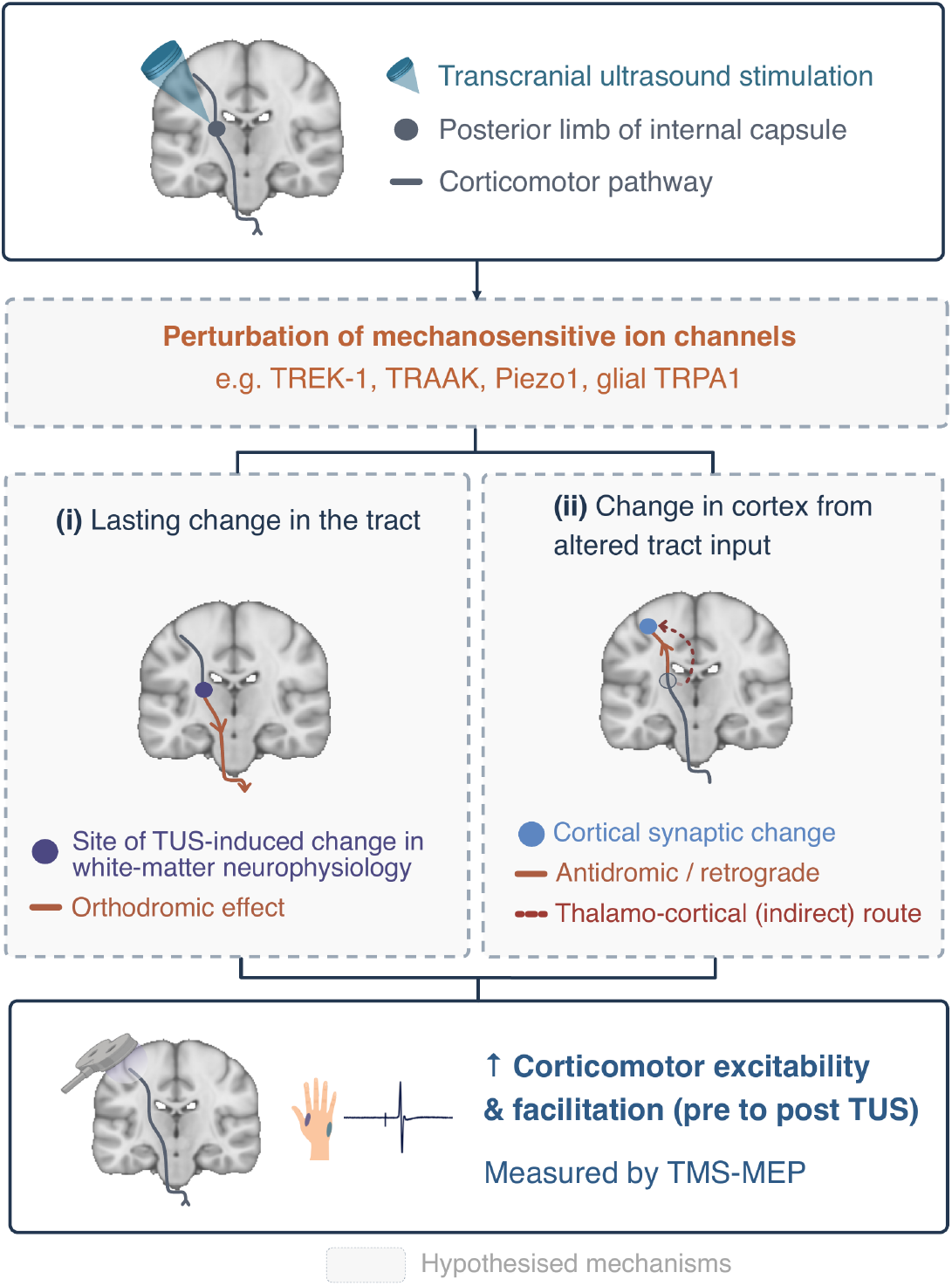
Hypothesised mechanisms of corticospinal tract TUS. TUS to the left posterior limb of the internal capsule increased corticomotor excitability and intracortical facilitation from pre- to post-sonication (TMS-MEP; solid boxes, observed). Intervening steps (dashed boxes) are hypothesised: perturbation of mechanosensitive ion channels could act via (i) a lasting change in the tract, altering the descending (orthodromic) corticomotor volley, or (ii) a transient tract change that drives lasting cortical synaptic change via antidromic/retrograde or thalamocortical routes, leaving the tract unchanged.

A consistent feature of our data was that ultrasound-induced increases in excitability were most pronounced at 50 minutes following TUS. This delayed trajectory is inconsistent with transient auditory or somatosensory confounds^61^, which would be expected to exert their strongest effects immediately after stimulation, and instead suggests an activity-dependent mechanism. Non-specific session-level factors, including expectancy, arousal, and repeated-measures drift^47^, are similarly unlikely to account for the effect: in our within-subject design, these would be expected to influence all conditions rather than the left CST condition specifically, and declining alertness across a long session would be more likely to reduce, rather than systematically increase, corticospinal excitability. The active comparison conditions were further matched for the sensory and procedural experience of sonication. This time course also offers mechanistic clues. Whereas prior TMS studies of motor cortex report peak effects within 5–20 minutes^20,24^, our later window resembles delayed effects reported in humans^19,62^, primate data^63^, and *in vitro* work^64,65^ showing progressive changes in neuronal excitability and synaptic transmission over hours. That both CST and M1 stimulation followed this slower trajectory suggests that ultrasound engages processes beyond immediate excitability shifts, on a timescale later than conventional NMDA (N-methyl-D-aspartate receptor)-dependent early-phase long-term potentiation (LTP) would predict. Characterising this temporal profile more precisely, including later windows up to 2– 4 hours, will be critical for understanding the mechanisms and durability of ultrasound-induced effects in humans.

Consistent with a previous study targeting M1^24^, the facilitation was carried by the SICF peaks (1.4 and 2.8 ms) rather than the intervening trough (2.2 ms), and was more robust at 1.4 ms. SICF peaks are attributed to interactions of excitatory indirect (I)-wave circuits and greater synchronicity of descending action potentials, whereas the troughs are sensitive to GABA_A_-mediated intracortical inhibition^66^. In contrast to past work, our changes in SICF were observed in the left CST as opposed to the left M1 sessions. Nonetheless, our work in combination with past data^24^ indicates that TUS may preferentially be sensitive to facilitation at I-wave peaks. Further work is needed to confirm this account.

### Fundamental studies involving stimulation of human white matter circuits

Our findings open new avenues for mechanistic tests of circuit function. Although white matter tracts are central to motor, cognitive, and affective processes^1,4,5^, methods to establish their causal role in humans remain scarce. TMS and transcranial electrical stimulation lack the depth and spatial precision to engage tracts reliably, whereas deep brain stimulation is highly precise but restricted to severe clinical indications. Although transcranial temporal interference stimulation has recently been shown to modulate deep brain regions^67,68^, its fields are diffuse, hindering clear attribution of effects to grey versus white matter. By demonstrating site-specific neuromodulation of a white matter tract with non-invasive focused ultrasound, we provide a framework for testing hypotheses about human white matter and its contribution to behaviour.

Our findings also have implications for using white matter as a control site in TUS studies. An important distinction to make in our data was that the physiological effects depended on stimulating the tract linked to our readout: targeting the left CST altered corticomotor measures, whereas sonication of the contralateral CST produced no detectable change. This highlights the need for careful tract selection in future studies if white matter is used as a TUS control site.

### Clinical applications of tract-targeted TUS

These findings support a potential therapeutic route for tract-targeted ultrasound. In neurological diseases such as stroke and spinal cord injury (see e.g., ref.^69^ for proof-of-concept of spinal cord TUS), targeting the CST with TUS may promote recovery by strengthening corticomotor output^70,71^. Notably, left CST stimulation increased excitability in both FDI and ADM, even though TMS targeting was optimised for FDI. This pattern is consistent with the dense, heterogeneous fibre organisation within the internal capsule and suggests that tract stimulation can facilitate distributed motor output across multiple muscles. Such effects could prove beneficial in clinical contexts, where restoring coordinated activation across muscles is more valuable than focal modulation alone.

In neuropsychiatry, white matter pathways are therapeutic targets. For example, deep brain stimulation of subgenual cingulate–centred white matter circuits can alleviate treatment-resistant depression^72,73^. Although we did not probe affective systems, the demonstration of site-specific neuromodulation of a white matter tract with non-invasive ultrasound provides a mechanistic basis to extend this approach to neural circuits implicated in psychiatric conditions. Consistent with this account, early TUS studies^27,28^ report feasibility with encouraging clinical outcomes. Our data extend on these previous findings by demonstrating how factors such as the tract microstructure can affect outcomes.

### M1 effect

Our M1 condition provided an opportunity to revisit early reports that repetitive 5 Hz TUS increased cortical excitability and reduced intracortical inhibition^20,24^. While these seminal findings inspired subsequent studies, replication efforts have yielded inconsistent outcomes, with some showing no change^47^ and others reporting opposite effects^25^, i.e. decreased excitation. Our results provide important context for this variability. Consistent in direction with these original reports, we observed increased corticomotor excitability and reduced intracortical inhibition, although the time course differed: Zeng et al. found effects maximal at 5 minutes that resolved by 60 minutes^20^, whereas our M1 effect emerged at 50 minutes following TUS. A key difference from past work^20,24,47^ is that we employed individualised acoustic modelling based on each participant’s skull anatomy to optimise sonication parameters and the TUS vector to maximise target engagement (see Supplementary Fig. S2), rather than relying on the TMS ‘hotspot’ on the scalp as a heuristic for transducer placement. Both our own (Fig. 5d) and others’ work^47^ indicates this is a poor proxy, which would lead to mistargeting of the ultrasound beam. Beyond targeting, we addressed other methodological concerns by verifying the fidelity of TUS transducer output in a hydrophone tank upon completion of data collection. Further, we monitored TMS coil stability throughout testing and report analyses that demonstrate exceptional accuracy and consistency. Together, these methodological refinements underscore the importance of targeting and validation in TUS research.

### Limitations and future directions

A limitation of our work was that we used a histological atlas to guide initial simulations. Incorporating individualised tractography for sonication planning would further improve precision, as in some participants, the left CST target tended to lie slightly (∼ 2 mm) posterior to the CST diffusion path. To address this limitation, we developed novel analytic approaches that characterised ultrasound–tract coupling through a composite engagement metric (overlap and alignment) and a microstructure–pressure interaction model. These exploratory analyses leveraged natural variability in targeting to examine how beam–tract geometry and tract microstructure relate to the magnitude of the physiological response. Such strategies provide a transferable framework for quantifying and interpreting individual differences in ultrasound targeting.

Another consideration is that we applied the same free-field intensity across participants. Tailoring *in situ* pressures on an individual basis may help standardise the intensity across conditions. Relatedly, we held the sonication protocol constant and varied only the target site, isolating site rather than acoustic parameters. Our 5 Hz pulse repetition frequency was adopted from parametric comparisons targeting the motor cortex^24^, but which parameters drive tract-targeted effects remains an important open question for future work.

Our design used active comparison sites rather than a sham condition, and we did not administer a formal condition-discrimination questionnaire, although subtle differences in perceptual experience during sonication would not be expected to influence the neurophysiological measures reported here. Incorporating an explicit assessment of perceptual discriminability between target sites would nonetheless be a useful refinement for future studies. Further, because the right CST session was always scheduled last, the left-versus-right CST comparison was not counterbalanced for session order, unlike the left CST versus left M1 comparison. Analyses restricted to the two counterbalanced sessions left these estimates unchanged and preserved the Session × Time interaction for CME-ADM, SICI and SICF, although the less sensitive two-session test did not reach significance for CME-FDI (Supplementary Materials). Finally, our sample size is modest. The primary contrast was within-subject and repeated across five time points, but a sensitivity analysis indicates the design was informative for effects of the magnitude of our primary contrasts but not for smaller ones (Methods), and the individual-difference analyses are correspondingly exploratory. Depending on the experimental design and *a priori* power analysis for the effect of interest, future TUS-TMS studies will likely require larger samples.

## Conclusion

Our findings provide causal evidence that transcranial ultrasound stimulation can modulate a human white matter tract. Targeting the CST produced a site-specific increase in corticomotor excitability and intracortical facilitation that exceeded both active comparison sites, the ipsilateral motor cortex and the homologous tract in the opposite hemisphere. Supporting analysis indicated that the magnitude of this effect depended jointly on ultrasound pressure and tract microstructure, underscoring the importance of both biological and biophysical factors. These results establish tract-targeted ultrasound as a tool for probing the causal role of white matter in human circuits and lay the groundwork for non-invasive neuromodulation of white matter tracts in disorders of white matter.

## Methods

### Participants

We recruited 18 healthy adults. Three participants did not complete the required sessions (two completed only the MRI session; one completed only one TUS-TMS session) due to scheduling constraints and were therefore not included in the analysed dataset, yielding a final sample of 15 participants (age = 30.87 ± 5.57 years; 5 female/10 male). As no prior data exist to inform a power calculation for our CST condition, we based our estimate on the effect size from a previous study targeting M1 (see Experiment 2 in ref.^20^), which indicated that 6 participants would provide 90% power (α = 0.05) to detect a within-subject interaction (G*Power 3.1.9.6). An *a priori* estimate derived from a single early report is approximate, and effect sizes in initial TUS-TMS studies have since proven variable across laboratories. We therefore characterise the sensitivity of the design rather than its power to detect the effect we observed. A parametric bootstrap over the design (1,000 simulations) indicated that between-session differences in linear trend of β ≈ 2.5 for FDI and β ≈ 1.2 for ADM would be detected with 80% probability. The design was therefore informative for effects of approximately the observed magnitude (FDI left versus right CST, β = 3.34; ADM, β = 1.61 and 1.31) but not for substantially smaller ones.

Exclusion criteria were age <18 or >50 years, recent use of psychotropic medication, a personal history of neurological disease or psychiatric illness, and contraindications to MRI, TMS or TUS^49,74^. All participants provided written informed consent. One participant did not complete the diffusion scan due to claustrophobia in the MRI machine limiting MRI session duration, but completed all other aspects of the study. The right CST control session was added after the left CST and left M1 sessions were complete, and 10 of the 15 participants attended. The experimental procedures were conducted in accordance with the Declaration of Helsinki and were approved by the Monash University Human Research Ethics Committee (ID #39671).

### Transcranial ultrasound stimulation (TUS) procedures

TUS was delivered using the NeuroFUS TPO system (Sonic Concepts Inc.) with a four-element annular transducer (CTX-500-101; diameter = 64 mm; central frequency = 500 kHz, steering range: 25–60 mm). Stimulation comprised a 5 Hz pulse repetition frequency with 20 ms pulse duration (pulse repetition interval = 200 ms), applied for 80 s (400 pulses total). The free-field spatial-peak pulse-average intensity (ISPPA) was fixed at 20 W/cm², measured in water before skull transmission.

To ensure safety, transcranial acoustic simulations were performed both *a priori* when planning for the TUS-TMS sessions and retrospectively after each session, confirming that intensities remained within the ITRUSST safety guidelines (MI and MI_tc_ ≤ 1.9)^49^. Thermal simulations were also conducted for the first 10 sessions, with maximum skull and brain temperatures remaining well below the 2°C threshold (mean = 0.4°C). In a self-report questionnaire completed 24 h after the final session, two participants reported a mild transient heating sensation during one of their TUS sessions (see Supplementary Materials).

To ensure reliable coupling, the hair was parted and centrifuged transmission gel (Aquasonic 100, Parker Laboratories) was applied, taking care to eliminate air bubbles^75^. A 10 mm gel pad (Hill Laboratories) was placed between the transducer and scalp. Neuronavigation was performed using Brainsight (v2.5.4, Rogue Research Inc.) and each participant’s T1-weighted structural image. During stimulation, the transducer trajectory was sampled every 5 seconds. The mean trajectory was used for confirmatory acoustic simulations, and positional error was calculated to verify that the transducer remained stable across the 80-second protocol (Fig. 5b). Bone conduction headphones played white noise at a constant volume in all sessions.

#### Target locations

The left CST target was defined at Montreal Neurological Institute (MNI) coordinates x = – 23, y = –20, z = 17, corresponding to a 100% probability region from the Jülich histological atlas (FSL). The right CST target (x = 25, y = –20, z = 17) was derived using the same process and atlas. CST targets were transformed from MNI to native space using FSL’s FLIRT and FNIRT for session planning and transducer placement. The M1 target was identified as the hand-knob region on the anterior bank of the central sulcus in each participant’s T1-weighted image, and further confirmed by our TMS ‘hot-spotting’ and motor mapping procedures.

#### Hydrophone calibration

We characterised the output of our TUS system using a 0.5 mm calibrated needle hydrophone (Precision Acoustics Ltd, Dorchester, UK) in a free-field setup. Measurements were acquired in a glass tank (81 × 36 × 40 cm) filled with degassed, deionised water (20.4 °C, total dissolved solids ≤ 1 ppm). The transducer was driven at a spatial peak pulse average intensity (ISPPA) of 20 W/cm² with the focal depth set to 55 mm, which corresponded to the average focal depth of our TUS-CST conditions. The hydrophone was mounted on a motorised 3D positioning stage, and line scans were performed along the axial and lateral axes of the ultrasound beam. Data acquisition and spatial control were implemented using MATLAB (R2023a, MathWorks, Inc.), with custom scripts adapted from an open-source hydrophone scanning framework (https://github.com/SamC873/FUSF_Hydrophone_Scanner). These measurements were used to validate the beam profile, i.e., the phase-controlled focusing of the four-element TUS transducer.

#### Acoustic simulations

Acoustic simulations were performed using the k-Wave Toolbox (v1.4) with custom MATLAB scripts (R2023a, MathWorks Inc.) on a high-performance computing cluster. Individual skull models were generated from each participant’s T1-weighted MRI using a deep-learning– based segmentation method^21,76^. We did not validate these pseudo-CT skull models against subject-specific CT for this scanner and sequence, and such MR-derived models remain difficult to validate in general^77^. Because simulated fields are sensitive to the assumed acoustic properties^77^, in-brain intensities should be treated as estimates carrying uncertainty^78^, which is a challenge for the field rather than one specific to our study. The simulated transducer was modelled according to the physical specifications of the NeuroFUS CTX-500 device. Simulations were run on a 256 × 256 × 256 grid centred on the midpoint between the transducer and acoustic focus (points per wavelength = 3).

### Transcranial magnetic stimulation (TMS)

Participants were seated with both hands resting pronated on a pillow in their laps. Surface electromyography (EMG) was recorded from the first dorsal interosseous (FDI) and abductor digiti minimi (ADM) of the right hand using Ag-AgCl electrodes (Ambu White Sensor WS, Ambu) in a belly–tendon montage. Signals were amplified 1000×, band-pass filtered at 10–1000 Hz (Digitimer D360 amplifier, Digitimer Ltd.), digitised at 10,000 Hz, and stored for offline analysis (Signal v6.05a, Cambridge Electronic Design). Each sweep began 100 ms before and ended 200 ms after the TMS trigger.

Monophasic TMS pulses were delivered over the left primary motor cortex using a D70 figure-of-eight coil (70 mm wing diameter) connected to two Magstim 200^2^ stimulators via a BiStim^2^ module (Magstim Co. Ltd). The FDI ‘hotspot’ was localised by identifying the hand knob region on the participant’s T1-weighted MRI and adjusting coil position under neuronavigation (Brainsight v2.5.4, Rogue Research Inc.) in 2 mm increments along the X (left–right) and Y (anterior–posterior) planes. Once identified, the trajectory was saved and held constant for the session (Fig. 5a). The coil was oriented tangentially to the scalp with the handle angled posteriorly at ∼45° to the central sulcus, inducing a posterior–anterior current flow in the brain. Resting motor threshold (RMT) was defined as the lowest intensity evoking MEPs ≥ 0.05 mV in 5 of 10 relaxed FDI trials. Active motor threshold (AMT) was defined as the lowest intensity evoking MEPs ≥ 0.2 mV during weak tonic contraction in 5 of 10 trials^74^.

Both single- and paired-pulse protocols were used (Fig. 1c–e). Corticomotor excitability (CME) was measured using a test stimulus adjusted to evoke ∼1.5 mV MEPs at baseline. Short-interval intracortical inhibition (SICI) was measured using a subthreshold conditioning stimulus 2 ms before the test stimulus, with the conditioning intensity set to produce ∼50% inhibition of the baseline MEP. In addition, we also collected SICI at 80% AMT (see Supplementary details and Fig. S7, confirming similar outcomes when using this approach). Short-interval intracortical facilitation (SICF) was measured by delivering the test stimulus followed by a second pulse at 100% RMT with inter-stimulus intervals (ISIs) of 1.4, 2.2, and 2.8 ms, corresponding to the first peak, trough, and second peak of the SICF time course^79^.

To ensure comparability of paired-pulse measures across time, test stimulus intensity was adjusted when necessary to maintain test MEP amplitudes within ± 30% of baseline. For all measures (CME, SICI, SICF_1.4_, SICF_2.2_, SICF_2.8_), 16 stimuli were delivered in pseudo-randomised order at 6 s intervals (±15% jitter; Signal v6.05a, Cambridge Electronic Design) at each time point. Each visit followed the same sequence: baseline TMS, TUS to the assigned target, and repeat TMS blocks at 5, 20, 35, and 50 minutes after the end of TUS.

### TMS-EMG analysis

All EMG recordings were screened offline, and trials with pre-stimulus muscle activity were discarded (root-mean-square ≥ 0.01 mV in the 100 ms before the TMS pulse). This resulted in the removal of 3.1% out of a total of 42,275 trials (2.9% for FDI; 3.3% for ADM). For single- and paired-pulse trials, motor evoked potentials (MEPs) were quantified as the peak-to-peak amplitude and averaged within each time point. For paired-pulse measures, conditioned MEP amplitudes were expressed as a ratio of the non-conditioned MEP (C/NC). To reduce the influence of outliers, an *a priori* trimmed-mean procedure was applied, discarding the highest and lowest value from each set of 16 MEPs^80^.

We analysed MEP amplitude for FDI and ADM, SICI ratios, SICF ratios at 1.4, 2.2, and 2.8 ms, and latency measures (reported in the Supplementary materials, Fig. S5) with separate linear mixed-effects models. Fixed effects included Time, Session, and their interaction, with participant as a random factor. Because corticomotor excitability, SICI, and SICF index mechanistically distinct cortical circuits rather than redundant tests of a single hypothesis^81^, we did not apply a family-wise correction across these measures. Multiplicity within SICF, which is assessed across several inter-stimulus intervals, was handled by modelling the inter-stimulus interval as a factor within a single model, and interval-resolved contrasts were Holm-corrected across the three intervals. Significant interactions were followed up with contrasts of model estimates and linear trends. Analyses were performed on non-normalised data for CME (normalised data are shown only for visualisation). Univariate outliers (1.3% of all data points), defined as values ± 3 standard deviations from the mean, were winsorised to one unit beyond the next most extreme value in the condition^82^. In one participant, two mid-session time points (20- and 35-min) in the left CST session were not acquired due to a brief interruption; in another participant, the FDI and ADM channels were only partially recorded due to a technical issue. These time points were treated as missing without imputation.

Linear mixed models were fitted using the *lme4* package^83^ in RStudio (v2023.06.0+421) with REML estimation, and Type III ANOVAs were conducted using Satterthwaite degrees of freedom approximation via *lmerTest*^84^. Linear time trends (slopes) were computed using centred time points using *emmeans*^85^, with between-session comparisons corrected using the Holm method. Time point contrasts were computed as model-based estimated marginal means using *emmeans*, and effect sizes were calculated as Cohen’s *d* for paired differences using the *effectsize* package^86^.

### TMS E-Field simulations (Fig. 1f–h)

The electric field (E-field) induced by TMS was quantified using SimNIBS 4.0^87^ integrated within the neuronavigation system (Brainsight v2.5.4, Rogue Research Inc.). For each participant and session, simulations were run using the coil position recorded during stimulation. Stimulation strength was converted from percentage of maximum stimulator output (% MSO) to device-specific dI/dt (A/s) using a custom MATLAB script calibrated for the Magstim D70 coil^88^. These values were entered into SimNIBS, and simulations were computed on subject-specific head models. Resulting E-field maps were transformed into MNI152 space, and group-level maps were obtained by averaging across participants.

### Motor mapping (Fig. 5c)

To verify that TMS ‘hotspot’ was optimised to the FDI representation within the hand area, we performed a motor mapping procedure for each participant. This was done by stimulating along the precentral gyrus from 10 mm lateral to 15 mm medial of the FDI ‘hotspot’. At each site, 5 pulses were delivered at 115% of resting motor threshold (RMT) at 6 s intervals (± 15% jitter) while recording MEP peak-to-peak amplitudes from the FDI and ADM. Stimulation sites were spaced at ∼3 mm increments, with positions determined by individual anatomy. This procedure confirmed the FDI representation at the identified ‘hotspot’, and indicated the ADM ‘hotspot’ was located 12.5 mm more medial, as expected (Fig. 5c).

### Magnetic resonance imaging (MRI)

MRI data were acquired with a Siemens 3T Skyra scanner and 32-channel head coil. We first acquired T1-weighted structural images (Magnetization Prepared Rapid Gradient Echo, TR = 2.3 s, TE = 2.07 ms, voxel size 1 mm^3^, flip angle 9°, 192 slices). Diffusion-weighted images were then acquired using a spin-echo EPI sequence (TR = 5900 ms, TE = 171 ms, voxel size = 2.5 mm isotropic, 56 slices, no gap) with 64 diffusion directions (b = 3000 s/mm²) and seven b0 images. The sequence employed monopolar gradients, GRAPPA acceleration (iPAT = 2), and whole-brain coverage with a field of view of 212 mm. A separate b0 scan with the opposite right-to-left phase encoding direction was acquired to enable susceptibility distortion correction.

### Diffusion-weighted imaging and tractography

Diffusion-weighted images were pre-processed in FSL, including susceptibility- and eddy current–induced distortion correction (*topup, eddy*) and brain extraction (*BET*) on the corrected b0 volume. Diffusion tensors were estimated with *dtifit*, and fibre orientation distributions were modelled using *bedpostx*.

T1-weighted anatomical images were registered to MNI152 standard space using FLIRT and FNIRT. ROIs corresponding to the posterior limb of the internal capsule (PLIC) and cerebral peduncle were derived from the Jülich probabilistic atlas (thresholded at 25%), inverse-warped into native space, and aligned to diffusion space using affine registration. The cortical seed was defined by the group-average TMS-induced E-field (thresholded at 40 V/m, which corresponds to the approximate intensity for resting motor threshold^89,90^) and projected into each participant’s diffusion space.

Tractography was performed using *probtrackx2* (10,000 streamlines per seed, step length = 0.5 mm, curvature threshold = 0.2), with the E-field derived cortical ROI as the seed, the PLIC as a waypoint, and the cerebral peduncle as the termination target. An exclusion mask of the contralateral hemisphere was applied to restrict streamlines to the ipsilateral CST. For each participant, tractograms were normalised by waytotal and thresholded at 1% of samples reaching the target to define individual CST masks. These were transformed into T1 and MNI space for downstream analysis and quantification of TUS–CST overlap.

### Probabilistic tractography analyses

#### Peak targeting (Fig. 4a)

To assess the spatial specificity of stimulation, we quantified ultrasound pressure within three regions defined in each participant’s anatomical T1 space. The target corticospinal tract (CST) was delineated using probabilistic tractography thresholded at ≥ 1% of streamlines reaching the target. The cortical stimulation zone was defined by SimNIBS-derived electric field maps thresholded at 40 V/m, and grey matter voxels were identified from SimNIBS tissue segmentation. Ultrasound pressures were summarised using the 99^th^ percentile to reduce outlier influence, and region-wise differences were tested with paired t-tests.

#### Tract engagement (Fig. 4b)

We quantified TUS targeting with a composite engagement index integrating (i) overlap between the ultrasound beam and the CST and (ii) alignment between the beam axis and the tract axis. Each component was z-scored after transformation, then summed:

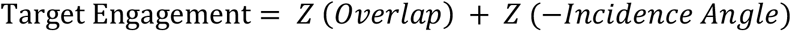

##### Overlap

Tract overlap was quantified by intersecting tractography-defined CST masks (≥ 1% of waytotal) with binarised acoustic pressure fields (> 0.25 MPa) in native T1 space. Overlap was computed as the percentage of CST voxels contained within the sonicated volume.

##### Incidence Angle

The beam axis was estimated via principal component analysis (PCA) on the coordinates of pressure voxels exceeding 0.25 MPa after projection to the sagittal (XZ) plane. The first principal component was normalised and oriented inferiorly (−Z) for consistency across participants. To characterise tract geometry at the beam–tract interface, we identified, for each participant, the peak-pressure voxel within the CST. A depth-restricted region (± 20 mm along the Z axis) centred on this voxel was intersected with the CST to isolate relevant tract voxels, which were then projected to the XZ plane. PCA on these coordinates yielded the dominant tract axis, which was normalised and oriented inferiorly. The incidence angle was defined as the acute angle between the unit beam and tract vectors:

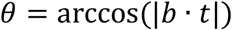

Where *b* and *t* are unit vectors of the beam and tract, respectively. Across participants, mean incidence angles were 27.5° ± 11.3° (range: 4.1° – 43.4°). The TUS-induced change in CME was defined as the mean change from baseline to the average of the 35- and 50-minute post-TUS time points for CME recorded from the FDI muscle.

#### White matter microstructure (Fig. 4c)

To assess whether microstructural properties modulated the effects of TUS, we extracted fractional anisotropy (FA) values at the site of ultrasound–tract interaction. For each participant, the peak pressure voxel was identified, and a depth-restricted analysis window (± 5 mm along the z-axis) was intersected with the tractography-defined CST to isolate relevant tract voxels. FA maps were registered from diffusion to T1 space, and mean FA was calculated within this tract-constrained mask. The relationship between FA, acoustic pressure, and changes in corticomotor excitability (CME) was modelled using least squares regression:

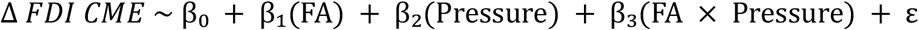

where Δ FDI CME was defined as the mean change from baseline to the average of the 35- and 50-minute post-TUS time points. The interaction term tested whether tract microstructure moderated the pressure–response relationship.

## Data & Code Availability

Data and analysis code will be made publicly available upon publication.

## Acknowledgements

D.C. is supported by a Monash University Early Career Postdoctoral Fellowship, a Monash University School of Psychological Sciences Strategic Project Grant, and a Marie Skłodowska-Curie Postdoctoral Fellowship (grant agreement no. 101280507). E.F.F. is funded by a UKRI FLF (MR/Y034368/1), a BBSRC (BB/Y001494/1) and an ARIA grant (SCNI-PR01-P15). J.J.H. is supported by an Australian Research Council Discovery Early Career Research Award (DE240101348). J.P.C. is supported by the Australian Research Council (FT230100656).

The authors acknowledge the facilities and scientific and technical assistance of the National Imaging Facility (NIF), a National Collaborative Research Infrastructure Strategy (NCRIS) capability at Monash Biomedical Imaging (MBI), a Technology Research Platform at Monash University. We acknowledge the technical assistance of Richard McIntyre. We also acknowledge Jayden Gagic for his work on the hydrophone tank used to characterise the transducer.

## Author Contributions Statement

*Conceptualisation*: D.C., J.J.H., J.P.C. *Formal analysis*: D.C., J.W., J.J.H., J.P.C. *Funding acquisition*: D.C., J.J.H., J.P.C. *Investigation*: D.C., J.W., J.J.H., J.P.C. *Methodology*: D.C., J.W., E.F.F., J.J.H., J.P.C. *Supervision*: E.F.F., J.J.H., J.P.C. *Visualisation*: D.C., J.J.H., J.P.C. *Writing – original draft*: D.C. *Writing – review & editing*: J.W., E.F.F., J.J.H., J.P.C.

