## Supplementary materials for "Transcranial ultrasound neuromodulation of a human white matter tract"

**Contents:**

**Fig. S1:** Individual Left CST targeting

**Fig. S2:** Individual M1 targeting

**Fig. S3:** Individual Right CST targeting

**Fig. S4:** Neuromodulatory and safety ultrasound characteristics

**Fig. S5:** MEP onset latency

**Table S1:** Transcranial magnetic stimulation (TMS) accuracy

**Table S2:** Transcranial ultrasound stimulation (TUS) accuracy

**Fig. S6:** Background muscle activity (rms EMG)

**Fig. S7:** 80% AMT SICI approach

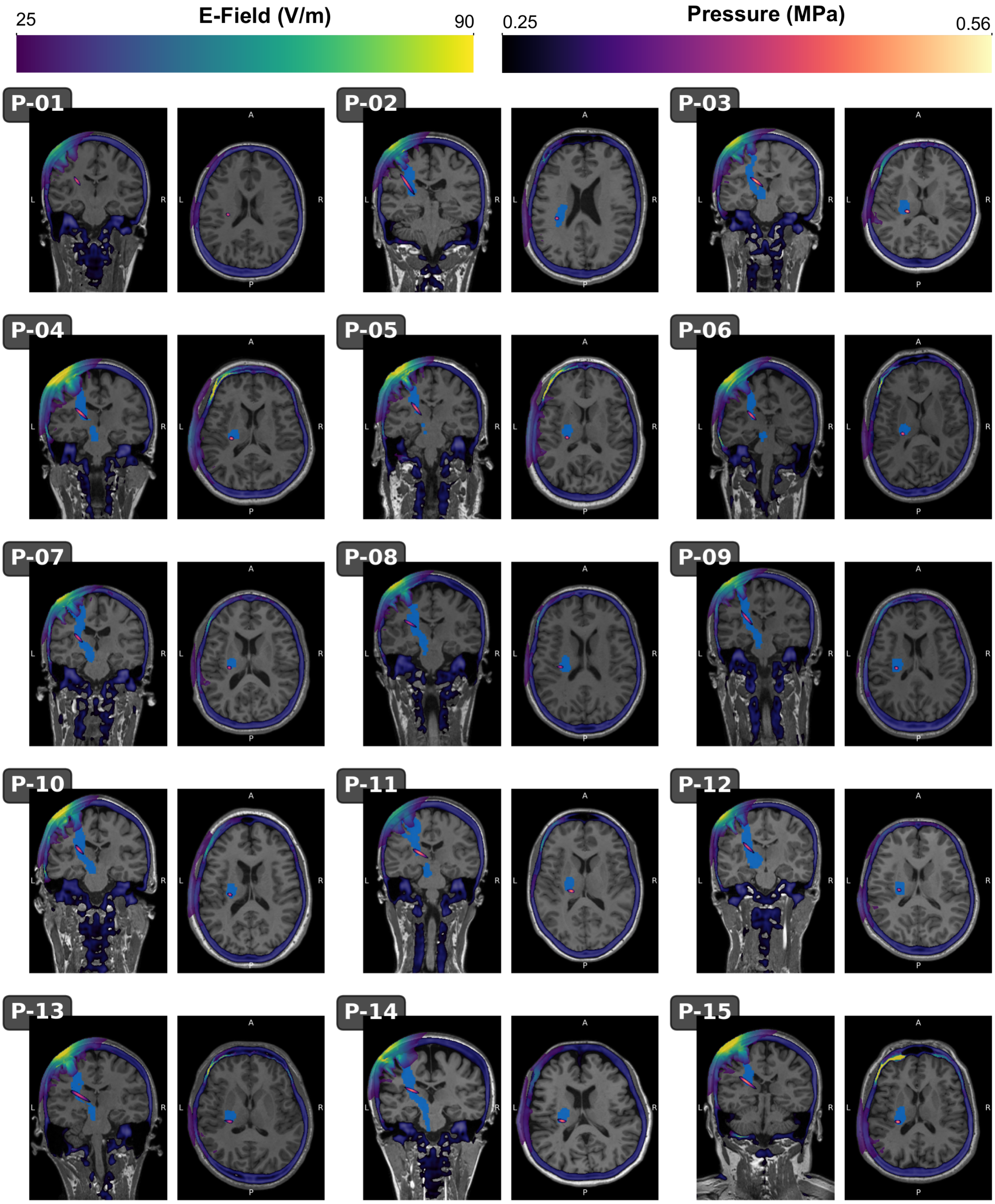


**Fig. S1. Individual Left CST targeting.** Individual coronal and axial slices in T1 anatomical space showing TMS electrical field (E-Field; *Viridis* colour scheme), 1% waytotal left CST tract (light blue transparent shading), pseudo-CT (thresholded at 300 HU; *Blue-Grey* colour scheme), and the acoustic pressure field (thresholded at 0.25 MPa; *Magma* colour scheme).

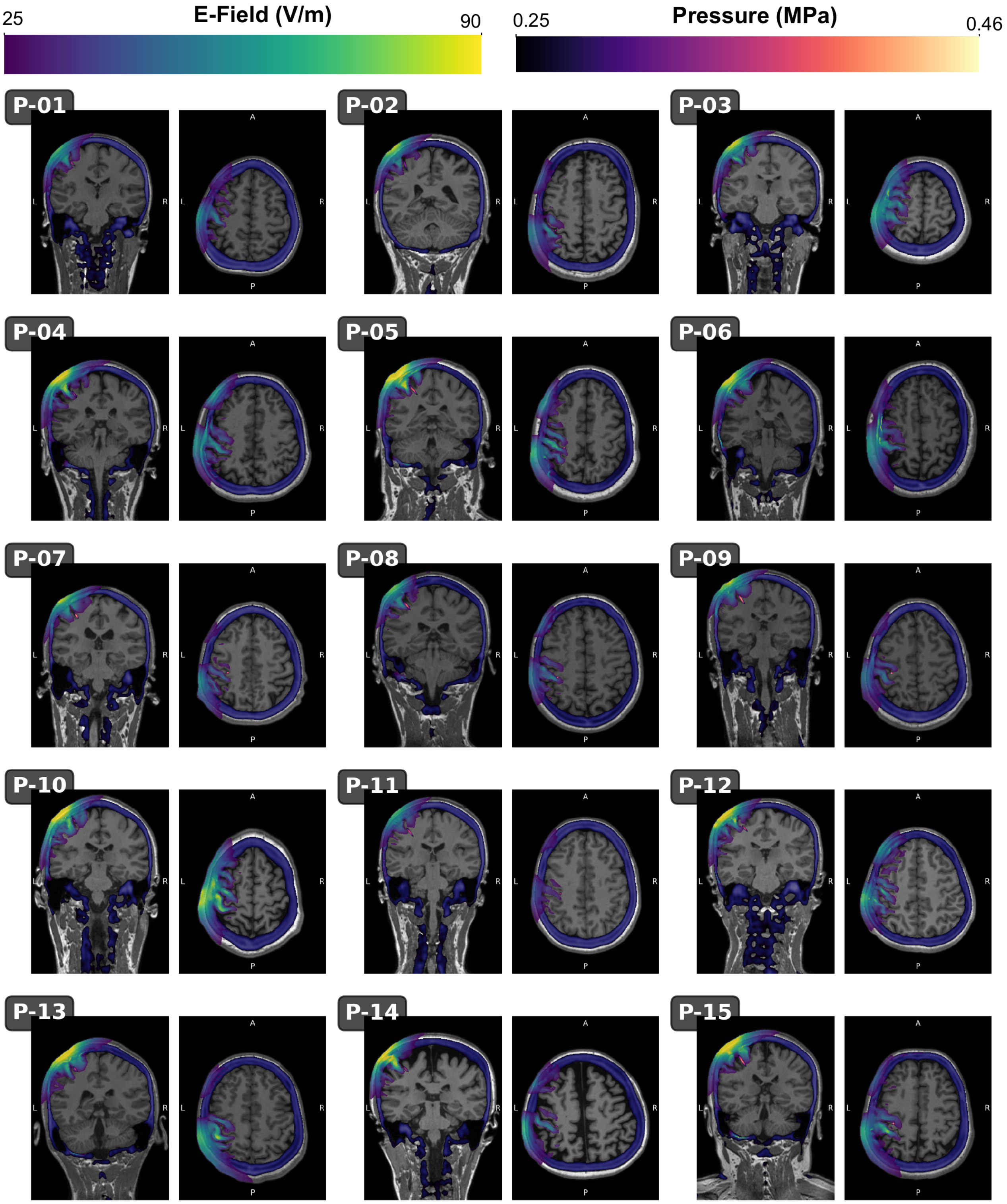


**Fig S2. Individual Left M1 targeting.** Individual coronal and axial slices showing TMS electrical field (E-Field; *Viridis* colour scheme), pseudo-CT (thresholded at 300 HU; *Blue-Grey* colour scheme), and the acoustic pressure field (thresholded at 0.25 MPa; *Magma* colour scheme).

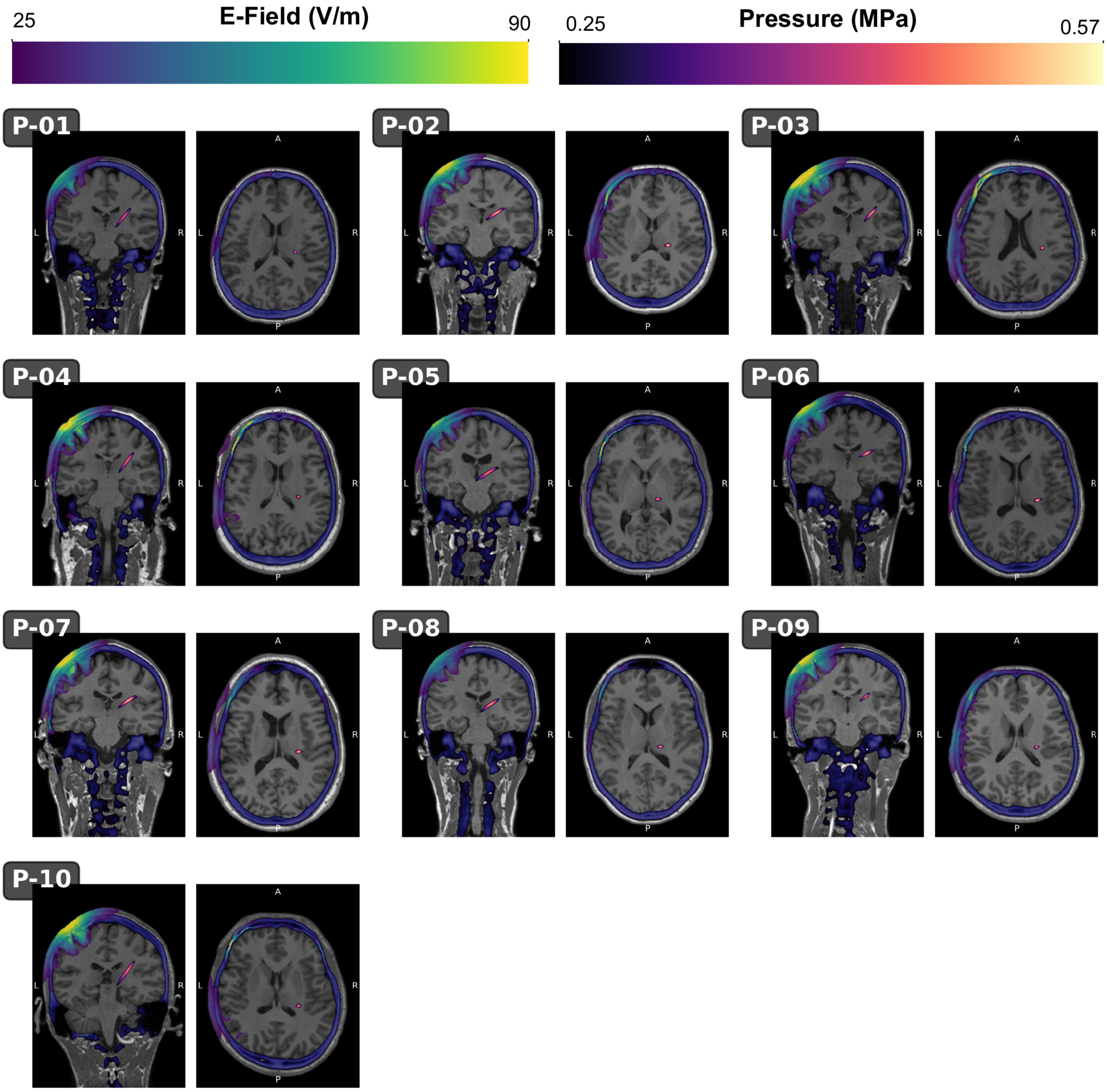


**Fig S3. Individual Right CST targeting.** Individual coronal and axial slices showing TMS electrical field (E-Field; *Viridis* colour scheme), pseudo-CT (thresholded at 300 HU; *Blue-Grey* colour scheme), and the acoustic pressure field (thresholded at 0.25 MPa; *Magma* colour scheme).

### Side Effects

A side-effects questionnaire was administered 24 hours after each participant's final session[1]. Participants rated potential side effects on a 4-point scale (absent, mild, moderate, severe) and provided open-ended responses describing any additional experiences and whether they attributed them to TMS or TUS. Two participants reported a mild heating sensation on the scalp during one of their TUS sessions. In one case, this lasted approximately 1 hour post-stimulation. Two participants, both with prior TMS exposure, reported *moderate* neck pain and numbness, which they attributed to TMS and prolonged sitting. No other side effects were reported.

### Neuromodulatory and safety ultrasound characteristics

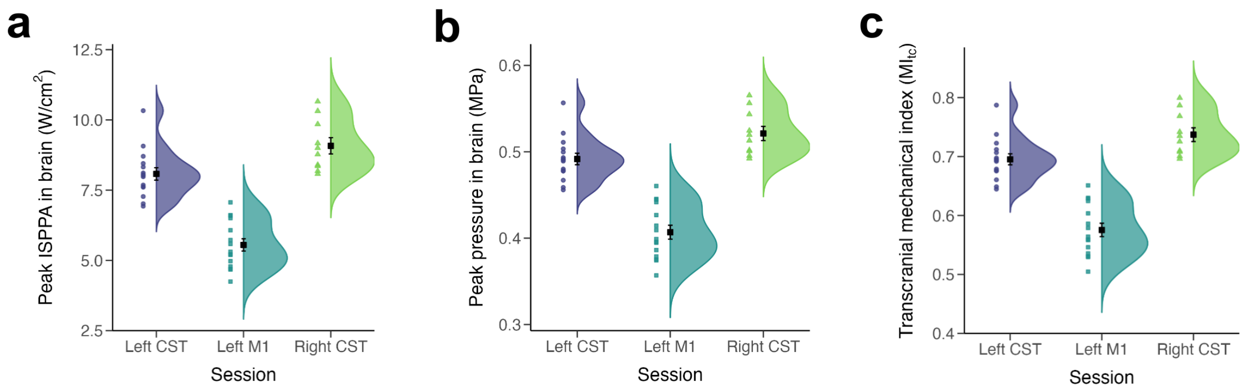


**Fig. S4. Neuromodulatory and safety ultrasound characteristics.** Average and distribution of peak ISPPA (W/cm²), peak pressure, and mechanical index, all within the brain. Data from each participant are presented on the left of each plot.

### MEP onset latency

Because ultrasound can alter the conduction of action potentials along myelinated axons[2], we tested whether TUS affected MEP latency. Here, latency was defined as the interval between single-pulse TMS delivery and onset of the muscle response. Latencies were determined using a custom semi-automated MATLAB script (MATLAB 2023a) that identified the first point at which the absolute EMG exceeded two standard deviations above the pre-stimulus baseline mean. All detections were verified manually and adjusted if necessary (< 5% of trials).

A linear mixed-effects analysis showed no main effects of Time or Session, and no interaction (all $p \geq 0.097$; Fig. S5). Given that the corticomotoneuronal pathway spans > 1 metre in most adults, while our sonication targeted only a short segment, any conduction-velocity change would be expected to produce sub-millisecond latency shifts, likely beneath the sensitivity of surface EMG. Direct tests of action potential conduction effects will therefore require more proximal methods, such as epidural spinal cord recordings[3].

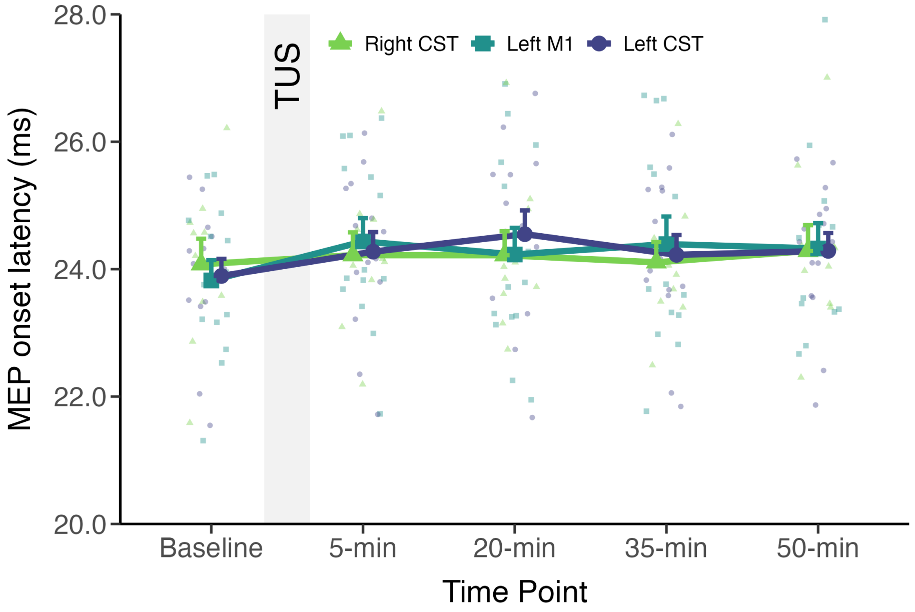


**Fig. S5. MEP onset latency.** Data represent the mean ± s.e.m. Individual participant data are presented as transparent data points.

### TMS and TUS consistency

Displacements in TMS coil position within a session, caused by participant or experimenter movement, could affect stimulation accuracy. To address this, we recorded coil position at every stimulation point and stored it in the neuronavigation system (Brainsight v2.5.4) for offline analysis. We then applied a drift-metric approach to quantify temporal changes in targeting accuracy across three measures: Target Error (mm), Angular Error (°), and Twist Error (°).

For each participant–session, two summary metrics were computed: (i) mean error, the average error magnitude across all stimulations, and (ii) slope error, the linear trend coefficient representing the rate of change in error across stimulations. For each error type and drift metric, we fitted the following linear mixed-effects models:

$$Error\ Metric \sim Session + (1 \mid Participant)$$

Session was a fixed effect with three levels (left CST, left M1, right CST), and Participant was a random intercept to account for individual differences.

**Table S1.** Transcranial magnetic stimulation (TMS) accuracy

| Error Type | Metric | Left CST | Left M1 | Right CST | Session |
|---|---|---|---|---|---|
| **Target Error (mm)** | Mean Error | 0.55 ± 0.15 | 0.55 ± 0.17 | 0.56 ± 0.21 | $F_{2, 26.31} = 0.05$, $p = 0.95$ |
| | Slope Error | 0.00013 ± 0.00042 | 0.00020 ± 0.00047 | 0.00015 ± 0.00035 | $F_{2, 28.20} = 0.13$, $p = 0.87$ |
| **Angular Error (°)** | Mean Error | 1.18 ± 0.58 | 1.18 ± 0.57 | 1.18 ± 0.74 | $F_{2, 25.35} = 0.00$, $p = 1.00$ |
| | Slope Error | -0.00036 ± 0.0013 | -0.00035 ± 0.0020 | -0.00018 ± 0.00082 | $F_{2, 27.65} = 0.14$, $p = 0.87$ |
| **Twist Error (°)** | Mean Error | -0.23 ± 0.40 | -0.17 ± 0.44 | -0.51 ± 0.39 | $F_{2, 27.42} = 3.00$, $p = 0.07$ |
| | Slope Error | 0.00073 ± 0.0011 | 0.00023 ± 0.0010 | 0.00075 ± 0.00089 | $F_{2, 40.00} = 1.18$, $p = 0.32$ |

Values are mean ± 1 standard deviation.

As an additional control, we recorded the TUS transducer position at 5-second intervals during each 80-second stimulation protocol. We applied the same drift-metric approach as for TMS, quantifying two error measures: Target Error (mm) and Angular Error (°). (Twist Error is not informative for TUS and was therefore excluded.) The same linear mixed-effects models were computed as described for TMS.

**Table S2.** Transcranial ultrasound stimulation (TUS) accuracy

| Error Type | Metric | Left CST | Left M1 | Right CST | Session |
|---|---|---|---|---|---|
| **Target Error (mm)** | Mean Error | 0.47 ± 0.27 | 0.54 ± 0.29 | 0.49 ± 0.21 | $F_{2,24.16} = 1.35$, $p = 0.28$ |
| | Slope Error | -0.011 ± 0.024 | -0.004 ± 0.028 | 0.008 ± 0.028 | $F_{2,39.00} = 1.59$, $p = 0.22$ |
| **Angular Error (°)** | Mean Error | 0.74 ± 0.56 | 0.66 ± 0.50 | 0.64 ± 0.31 | $F_{2,22.64} = 0.20$, $p = 0.82$ |
| | Slope Error | -0.002 ± 0.038 | -0.004 ± 0.035 | -0.022 ± 0.030 | $F_{2,37.00} = 1.12$, $p = 0.34$ |

Values are mean ± 1 standard deviation.

### TMS vs TUS coil position (Fig. 5d)

To quantify differences in transcranial ultrasound stimulation (TUS) targeting relative to transcranial magnetic stimulation (TMS) at the level of the scalp, we calculated positional offsets between TUS and TMS trajectory coordinates for the left primary motor cortex (M1) target. For each participant, 2D Euclidean distances (x–y plane) were computed, providing measures of targeting accuracy in the coronal plane. Spatial distributions were visualised using scatter plots with a *viridis* colourmap (where higher distances = lighter colours) colour-coded by 2D distance magnitude. The reference TMS position was fixed at the coordinate origin (0,0), with a 15 mm gradient circle indicating the approximate extent of the TMS electric field. For TUS, 5 mm diameter gradient circles were included around each data point to represent the average full width at half maximum. Ridgeline density plots show the distribution of positional differences along each anatomical axis. On average, the mean 2D targeting difference was 7.38 ± 3.97 mm (mean ± s.d.; range 1.37–16.92 mm), demonstrating the spatial offset between TMS and TUS targeting approaches for motor cortex stimulation at the level of the scalp.

### Assessment of order effects

To assess potential order effects, we fitted linear mixed models testing Session Number × Time Point interactions for each outcome, restricted to the left CST and left M1 sessions. Models included random intercepts for participant. No interaction effects were observed for CME (FDI: $p = 0.316$; ADM: $p = 0.819$), SICI ($p = 0.475$), or SICF at 1.4 ms or 2.8 ms ($p =$

0.141 and 0.126, respectively). A single order effect was detected for SICF at 2.2 ms ($p$ = 0.041), but this did not affect the interpretation of the main findings, as the primary effects for this measure were non-significant.

Because the right CST session was always scheduled third, session order carries no variance in that condition and cannot be modelled for it. We therefore also refitted each model using only the two counterbalanced sessions, left CST and left M1, in which session order was pseudo-randomised. Estimates were essentially unchanged from the models reported in the main text. Session × Time interactions were preserved for CME-ADM ($F_{4, 124}$ = 3.98, $p$ = 0.005), SICI ($F_{4, 120}$ = 3.30, $p$ = 0.013) and SICF ($F_{4, 383.52}$ = 2.74, $p$ = 0.029). Left CST differed from left M1 in the linear trend for CME-ADM (β = 1.31, 95% CI [0.54, 2.09], $t_{124}$ = 3.35, $p$ = 0.001, $d$ = 0.90) and SICI (β = −0.87, 95% CI [−1.40, −0.34], $t_{120}$ = −3.23, $p$ = 0.002, $d$ = 0.66), and at 50 minutes for CME-ADM (Δ = 0.48, 95% CI [0.23, 0.72], $t_{124}$ = 3.88, $p < 0.001$, $d$ = 0.68), SICI (Δ = −0.38, 95% CI [−0.55, −0.22], $t_{121}$ = −4.51, $p < 0.001$, $d$ = 0.75) and SICF at 1.4 ms (Δ = 0.96, 95% CI [0.27, 1.64], $t_{384}$ = 2.74, $p$ = 0.006, $d$ = 0.31). For CME-FDI the estimate and effect size were identical to the full model (β = 1.69, $d$ = 0.53), with $p$ = 0.061 in the reduced model. The two-session interaction is a less sensitive test than the three-session interaction reported in the main text, as it omits the condition in which the response differed most, and for CME-FDI it did not reach significance ($F_{4, 120}$ = 1.08, $p$ = 0.371). Differences were confined to the later time points, with none at 5 or 20 minutes for any measure (all $p > 0.25$), and the SICF effect remained largest at the 1.4 ms I-wave peak, with no change at the 2.2 ms trough.

**Ultrasound intensity differences**

We tested whether between-session differences in ultrasound intensity had an impact on the observed effects. For each measure, we computed a change score (Δ = 50-min post-TUS – baseline) and fit mixed models with Session and maximum simulated in-brain ISPPA as predictors. Maximum ISPPA (W/cm²) did not predict Δ for CME (FDI: $p$ = 0.223; ADM: $p$ = 0.441) or SICI ($p$ = 0.435), and session effects remained (FDI: $p$ = 0.007; ADM: $p < 0.001$; SICI: $p$ = 0.044). For SICF, neither intensity nor session significantly predicted Δ at any ISI (1.4 ms: intensity $p$ = 0.221, session $p$ = 0.102; 2.2 ms: intensity $p$ = 0.397, session $p$ = 0.486; 2.8 ms: intensity $p$ = 0.578, session $p$ = 0.150).

### Background rmsEMG

To verify that subthreshold muscle activity (≤ 0.010 mV) immediately prior to each TMS pulse did not influence our results, baseline rmsEMG (root-mean-square of the 100 ms before the TMS pulse) was analysed using linear mixed-effects models for each TMS measure (CME FDI, CME ADM, SICI, $SICF_{1.4}$, $SICF_{2.2}$, $SICF_{2.8}$).

$$rmsEMG \sim Session \times Time + (1 \mid Participant)$$

Although some session main effects reached statistical significance, these corresponded to negligible differences (0.0004–0.001 mV) that are not physiologically meaningful. Critically, there were no Session × Time interactions (all $p > 0.32$) and no main effects of Time (all $p > 0.32$), confirming that baseline rmsEMG activity remained stable across sessions and time points (Fig. S6).

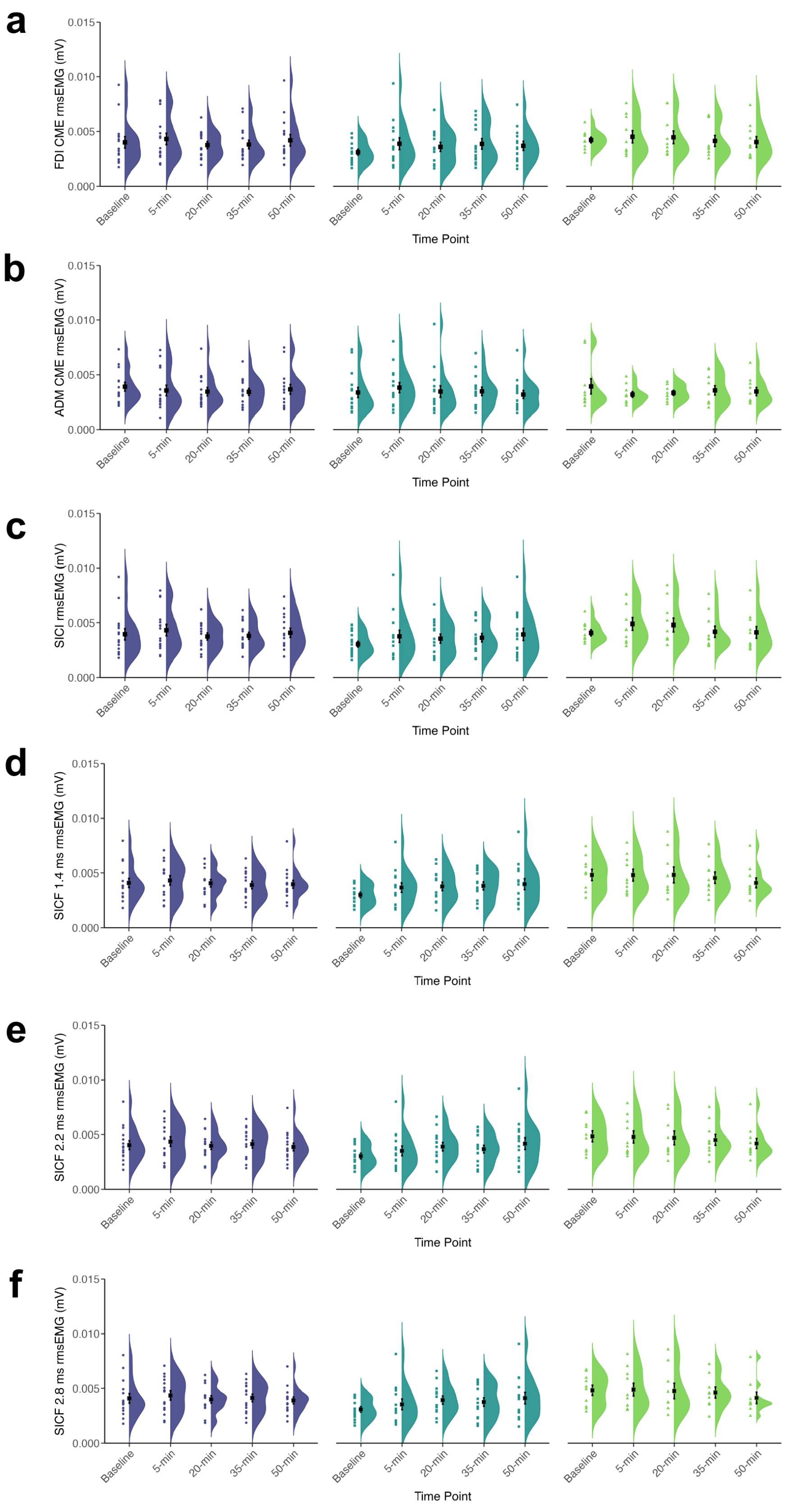


**Fig. S6. Background muscle activity.** Pre-trigger EMG activity showed no main effect of Time and no Session × Time interaction for any measure (all $p > 0.32$). Session differences reached significance for some measures but corresponded to negligible magnitudes (0.0004 to 0.001 mV). RMS = root mean square, EMG = electromyography, CME = corticomotor excitability, SICI = short-interval intracortical inhibition, SICF = short-interval intracortical facilitation.

### SICI CS50 vs 80% AMT analysis

We collected SICI in two ways: (i) with the conditioning stimulus set to 80% of AMT, and (ii) by adjusting the stimulator output to achieve approximately 50% inhibition. In many cases, the latter adjustment was unnecessary, as 80% AMT typically corresponds to ~ 50% inhibition. When required, we prioritised the CS50 approach in the main manuscript, as it is sensitive to both increases and decreases in inhibition. To assess whether conditioning intensity affected outcomes, we also analysed SICI collected at 80% AMT.

Using the same linear mixed-effects models, we confirmed that both approaches yielded a similar pattern of results (see Fig. S7), the same main effects of Session ($F_{2, 167.35} = 4.69$, $p = 0.01$) and Time ($F_{4, 164.82} = 3.51$, $p = 0.009$), but no Session × Time interaction ($F_{8, 164.82} = 1.64$, $p = 0.12$).

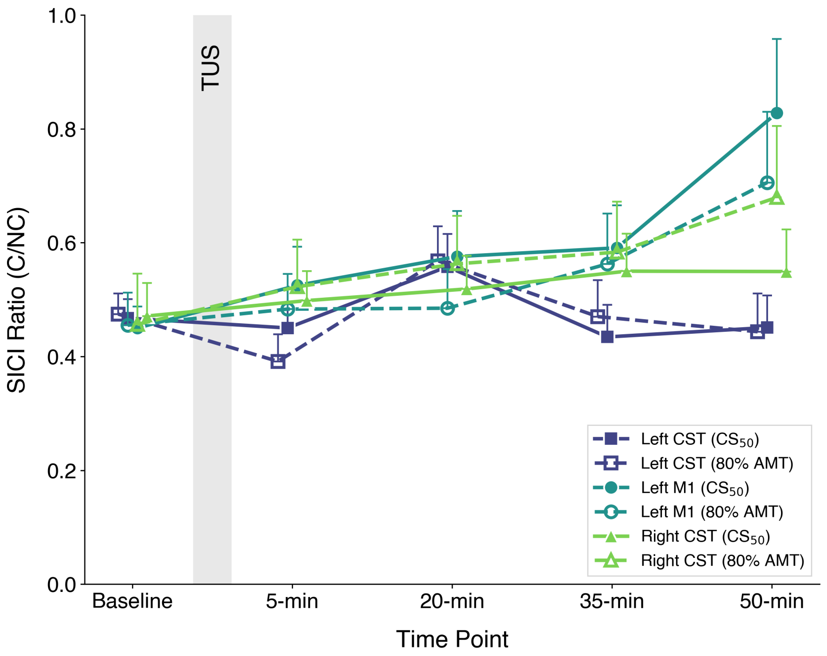


**Fig. S7. 80% AMT SICI approach.** Short-interval intracortical inhibition (SICI) ratios across the left corticospinal tract (Left CST), left primary motor cortex (Left M1), and right corticospinal tract (Right CST) conditions. The SICI ratio is expressed as the conditioned/unconditioned MEP amplitude.
